# A broad matrix metalloproteinase inhibitor with designed loop extension exhibits ultrahigh specificity for MMP-14

**DOI:** 10.1101/2022.12.29.522231

**Authors:** Alessandro Bonadio, Bernhard L. Wenig, Alexandra Hockla, Evette S. Radisky, Julia M. Shifman

## Abstract

Matrix metalloproteinases (MMPs) are key drivers of various diseases, including cancer. While several antibodies against MMPs are in development, our goal is to construct therapeutic anti-MMP inhibitors based on a natural broad MMP inhibitor, tissue inhibitor of metalloproteinases-2 (N-TIMP2). To confer high binding specificity toward one MMP type, we extend one of the N-TIMP2 loops, allowing it to interact with the non-conserved MMP surface. Multiple computational designs of the loop were used to design a focused library for yeast surface display, which was sorted for high binding to the target MMP-14 and low binding to off-target MMP-3. Deep sequencing of the two selected populations followed by comparative data analysis was used to identify the most promising variants, which were expressed, purified, and tested for inhibition of MMP-14 and off-target MMPs. Our best N-TIMP2 variant exhibited 29 pM binding affinity to MMP-14 and 2.4 µM affinity to MMP-3, 7500-fold more specific than WT N-TIMP2. Furthermore, the variant inhibited cell invasion with increased potency relative to WT N-TIMP2 in two breast cancer cell lines. We obtained the engineered variant high-accuracy model by including NGS data as input to AlphaFold multiple sequence alignment (MSA). Modeling results together with experimental mutagenesis demonstrate that the loop packs tightly against non-conserved residues on MMP-14 and clashes with MMP-3. This study demonstrates that introduction of loop extensions into inhibitors to stretch to the non-conserved surface of the target proteins is an attractive strategy for conferring high binding specificity in design of MMP inhibitors and other therapeutic proteins.

## Introduction

Matrix metalloproteinases (MMP) are a family of proteases, comprising twenty-three different family members in humans. They are composed of multiple domains, including a conserved catalytic domain with a catalytic zinc ion, an inhibitory pro-domain, and other domains (1). MMPs are involved in multiple biological processes, including remodeling of the extracellular matrix (ECM), shedding of cell surface receptors and membrane-bound signaling molecules, angiogenesis, intravasation/extravasation from blood vessels, and immune cells maturation (2). It is not surprising that MMPs play an important role in cancer metastasis, with multiple MMPs overexpressed in solid tumors (3–6). Many MMPs are associated with a more aggressive metastasis and tumor proliferation with MMP-9 and MMP-14 having a major role, while other MMPs like MMP-3 and MMP-8 have been reported to exhibit anti-tumor effects (2, 4, 7).

Due to the important role of MMPs in cancer, there has been considerable effort to develop therapeutics against these targets. Zinc chelators were the first small molecule inhibitors developed against MMPs (8). These inhibitors bind to the catalytic zinc in the MMP active site, which is highly conserved among all MMP family members as well as other similar enzymes. Due to their low binding specificity, they exhibited high toxicity and were not pursued as therapeutics (9). Subsequently, other small molecule inhibitors have been developed, yet directing them toward specific MMP family members remains a challenge (9– 11).

Unlike small molecules, proteins have the potential to bind large protein surfaces and hence to exhibit high binding specificity for a particular target protein (12). Thus, most of the latest efforts for MMP drug design have been focused on protein engineering. Several antibodies have been developed against various MMP targets with most efforts focusing on MMP-9 and MMP-14 (13–21). The monoclonal antibody Andecaliximab specifically binds MMP-9, at the junction between the catalytic domain and the pro-domain, and is being evaluated in multiple phase 3 clinical trials involving solid tumors (15, 16). Specific inhibitory anti-MMP-14 antibodies have been discovered through phage display technology (17), by using a library with long CDR-H3 to facilitate the binding to the enzyme catalytic cleft (18) and by using as immunogen a mimic of a loop far from the active site (19, 20).

While several anti-MMP antibodies are currently being evaluated in clinical trials and preclinical research, we have focused our effort on engineering specific MMP inhibitors starting from a broad family MMP inhibitor, tissue inhibitor of metalloproteinases-2 (TIMP2). TIMP2 is one of four homologous endogenous MMP inhibitors and it interacts with the conserved MMP active site as well as with some less conserved neighboring regions and inhibits all MMP family members with similar affinities (22). The N-terminal domain, N-TIMP2, is frequently used for protein engineering since it is a small 127 amino acids protein scaffold, stabilized by disulfide bonds (23, 24). N-TIMP2 retains low nM affinity to MMPs, moreover unlike the full-length TIMP2, it is not involved in MMP activation (25, 26). In addition, as an endogenous protein, N-TIMP2 exhibits no toxicity to humans. Furthermore, the potential of TIMP2 as a therapeutic has been recently demonstrated in a study where TIMP2 showed suppression of both primary and metastatic tumor growth in a murine model of triple-negative breast cancer (27).

We and others have engineered N-TIMP2 with the goal of converting it into a highly specific inhibitor of one MMP type (28–32). Our group first explored how various single mutants in N-TIMP2 affect its affinity and specificity to various MMPs (29). We subsequently used computational analysis to design focused libraries of N-TIMP2, where a small set of binding interface positions on the N-TIMP2 was randomized and such libraries were selected for binding to MMP-14 and MMP-9 with yeast display technology (30). The engineered N-TIMP2 variants exhibited high affinity and enhanced specificity toward MMP-14 and/or MMP-9 and significantly inhibited invasion of the cancer cells compared to the negative control, demonstrating great potential for future therapeutics development. However, the majority of previous N-TIMP2 designs demonstrated improved affinities to off-target MMPs, thus showing only moderate increase in binding specificity (31).

In the current study, we present a novel strategy for enhancing N-TIMP2 binding specificity through loop extension. Our strategy takes an advantage of the fact that although MMPs are very conserved at or near the active site, this conservation decreases at locations further from the active site (Figure 1A). Hence we thought to extend one loop of N-TIMP2 to enable its interaction with these MMP regions of high diversity, supplying it with specificity function found in natural loops (33). Protein loops have been previously engineered for increased affinity and specificity using directed evolution (34, 35) or computational design (36–39). In these studies, highly specific engineered loops exhibited backbone conformations preorganized for interactions and buried hydrogen bonds and charges upon complex formation (40).

**Figure 1.**
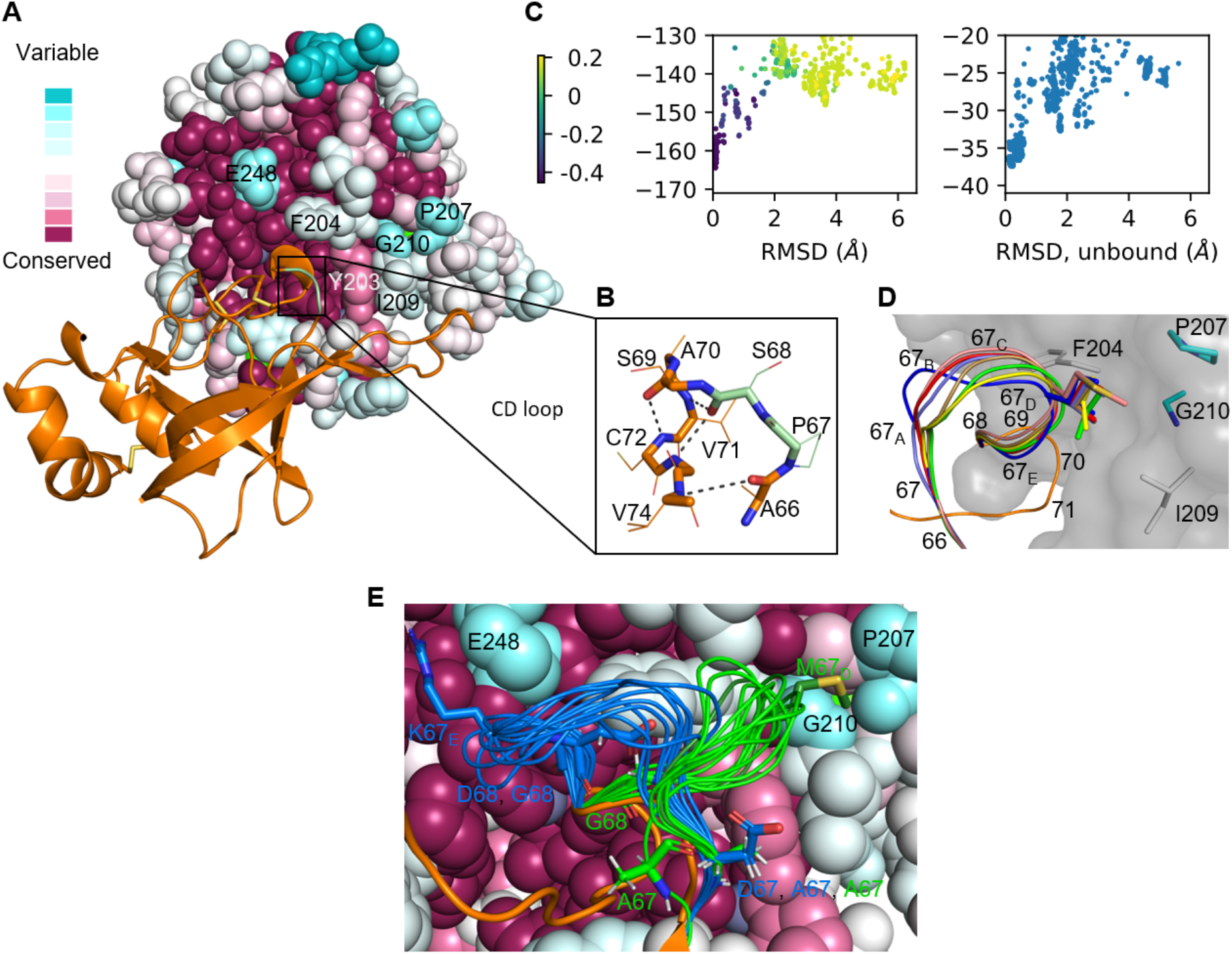
Design of an extended N-TIMP2 CD loop to reach non-conserved MMP regions. A, N-TIMP2/MMP-14 complex structure showing MMP conservation profile. N-TIMP2 is shown in orange and MMP-14 is shown in sphere representation (PDB ID: 1BUV (42)) colored by sequence conservation calculated with the ConSurf web server (41). The N-TIMP2 CD-loop is shown in green. B, zoom into the CD loop. A66 forms a backbone/backbone hydrogen bond with V74 and S69 forms a backbone/backbone hydrogen bond with C72. C72 is further anchored by a disulfide bond with C1. The engineered loop comprises a 7-residue insertion between the anchor residues A66 and S69. C, sampling of the loop conformation of a design (Des4) by Rosetta FKIC in the MMP-14-bound (left) and unbound (right) states. The lowest energy models are very similar to the design model both by RMSD of the loop and target residue binding energy (dark purple), in the bound and unbound conformations. The color of the dots on the left figure corresponds to the sum of binding energy to the target residues. D, lowest energy conformations of the seven selected designs, showing loop position 67_D_ interacting with target residues on MMP-14. E, two clusters of sequence designs from an unrestrained loop design contacting target residues on MMP-14.

Here, we computationally extended the CD loop on N-TIMP2 to reach the non-conserved MMP regions and designed a combinatorial library of N-TIMP2 variants. We then utilized yeast surface display (YSD) coupled to next generation sequencing (NGS) to select N-TIMP2 loop variants that bind with high affinity to the target MMP-14 and with low affinity to the off-target MMP-3. Several engineered N-TIMP2 mutants were experimentally characterized, with the best variant exhibiting 83000-fold binding specificity toward MMP-14 relative to off-target MMP-3, 7500-fold improvement in specificity compared to WT N-TIMP2. The variant inhibited breast cancer cell invasion with increased potency relative to WT N-TIMP2.

## Results

### Design of N-TIMP2 variants with extended CD-loops

To design more specific N-TIMP2 variants by loop extension, we first analyzed MMP conservation using the ConSurf web server (41). The conservation pattern shows that the region at and close to the MMP active site is highly conserved, while more distant residues show medium and low conservation profiles (Fig. 1A).

N-TIMP2 interacts with MMP-14 through the N-terminus and three loops with two of them binding to the highly conserved parts of MMP-14 and the third one binding to a less conserved region (Figure 1A). Inspection of the N-TIMP2 loops revealed that it would be possible to remodel the CD loop of N-TIMP2 to reach the generally non-conserved MMP region, without affecting the overall N-TIMP2 fold. The newly generated N-TIMP2/MMP-14 interactions could result in enhanced affinity of N-TIMP2 to MMP-14 and simultaneous decrease of affinity to off-target MMPs due to the introduction of apparent clashes and non-optimal interactions.

We next examined various points for loop insertion on the CD loop considering that insertion points should be well anchored in the N-TIMP2 structure to preserve the protein fold and to make the loop rigid. We initially identified T65 and V71 as the points for loop insertion, reasoning that T65 is the last amino acid of the β-strand and as such is well anchored in the structure, while V71 precedes C72, which is anchored by a disulfide bond (Fig. 1B). We then computationally designed loops of different lengths to fill the gap between residues T65 and V71 on N-TIMP2, inserting 8 to 11 residues and applying restraints to force the loop to interact with the target residues on MMP-14. Visual inspection showed that many models recapitulated the structure of the WT CD loop at the first N-terminal amino acid (A66) and the last two C-terminal amino acids (S69 and A70), which are indeed anchored to the protein by backbone/backbone hydrogen bonds. To increase the accuracy of the loop design, we decided to keep A66, S69, and A70 as WT and to introduce a 7-amino-acid loop insertion between residues A66 and S69 (Fig. 1B).

We next constructed 2000 CD loop models in the context of the N-TIMP2/MMP-14 structure using Rosetta Remodel (37), with a *de novo* loop building protocol that assembles loop fragments and subsequently closes the loop. 35 designs that showed the best interaction energy with two non-conserved target residues on MMP-14 and the best total energy were selected and the loop sequence was redesigned using Rosetta Relax (43). We then performed loop modeling of the designed sequences in Rosetta using kinematic closure with fragments (FKIC)(44)(45), modeling both the bound (N-TIMP2/MMP-14) and the unbound (N-TIMP2 alone) conformations (Fig. 1C, Supplementary Fig. S1). Designs were analyzed to determine whether the designed loops exhibited a single low-energy state as such properties would be beneficial for enhancing binding specificity. Seven designed loop sequences were selected based on these criteria (Fig. 1D, Supplementary Table S1).

While designing protein binders typically requires multiple cycles of design and testing with sometimes tens of thousands of designs being evaluated (12, 46, 47), in the current work we decided to increase the success of the loop designs by constructing a focused library of N-TIMP2 variants where loop randomization was based on design calculations. In the library design, we allowed only two consensus amino acids at positions 67 and 68 since these positions were shown to be more spatially restricted compared to the rest of the loop in multiple designed loop clusters and the seven selected designs (Table 1, Fig. 1D, Fig. 1E, Supplementary Fig. S2). We fully randomized positions 67_D_ and 67_E_ since they were at the interface with MMP-14 in all designed loop clusters and the 7 selected designs; the remaining three positions were soft randomized based on the sequences of the clusters and the seven selected designs, prioritizing the seven selected designs. The loop-neighboring positions 66 and 71 were not mutated in the modeling protocol but were randomized in the library to allow for the introduction of loop-stabilizing interactions and relaxation of possible clashes. The resulting library contained 3.1*10^8^ variants, about two thousand-fold smaller than a 9-residues fully randomized library (5.1*10^11^), enabling exhaustive testing of this library by YSD.

**Table 1.** Design of focused CD loop library for N-TIMP2.

|  | **66** | **67** | **67<sub>A</sub>** | **67<sub>B</sub>** | **67<sub>C</sub>** | **67<sub>D</sub>** | **67<sub>E</sub>** | **68** | **71** |
| --- | --- | --- | --- | --- | --- | --- | --- | --- | --- |
| WT | A | P | - | - | - | - | - | S | V |
| Library | all | AD | ADEIKMNTV | ADEGKNRST | ADGNST | all | all | DG | all |
| # aa | 20 | 2 | 9 | 9 | 6 | 20 | 20 | 2 | 20 |

### Selection of N-TIMP2 variants for binding to MMP-14

We used YSD to select N-TIMP2 variants with high-affinity binding to MMP-14. In the YSD setup, N-TIMP2 was displayed on the surface of the yeast cells with a free N-terminus and a C-terminal Myc-tag (Fig. 2A) as leaving the N-terminus free is critical for binding to MMPs. N-TIMP2 expression was detected using phycoerythrin (PE) conjugated anti-Myc tag antibody while MMP binding was detected using DyLight-488 labeled MMPs. The designed library of N-TIMP2 loop variants was experimentally constructed and transformed into yeast yielding 1.9*10^8^ transformants.

**Figure 2.**
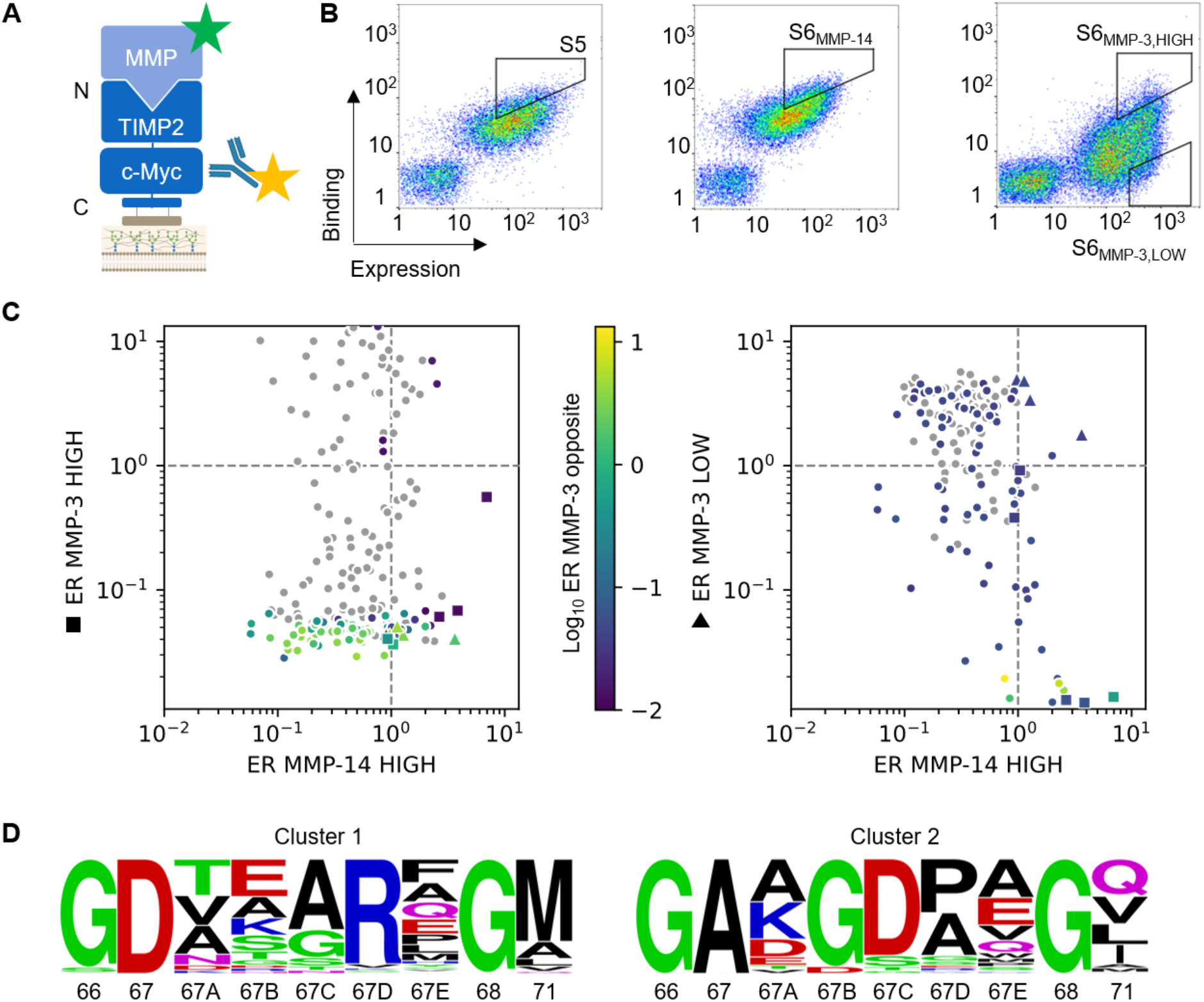
Identification of selective variants using YSD and deep sequencing. A, the YSD setup for N-TIMP2 variant selection. A library of N-TIMP2 variants was synthesized and transformed into yeast and displayed on the surface of yeast cells. Yeast cells were incubated with a PE-conjugated α-Myc tag antibody, allowing monitoring of N-TIMP2 variant expression, while MMP-14 was labeled with Dylight-488 and incubated with the yeast cells. B, the last three YSD selections that were subjected to NGS. Selection gates are shown and labeled with the sorting cycle and the MMP used. For S5 and S6_MMP-14_, 20nM MMP-14 was used, and for S6_MMP-3,HIGH_ and S6_MMP-3,LOW_, 1 µM MMP-3 was used. Diagonal gates were set to collect the portion of the population exhibiting more than half the maximum or the lowest binding/expression signal. C, correlation between enrichment ratios (ER) in sorts for the positive target and for the negative target with high (left) and low (right) binding affinities for a few specific variants identified by NGS. Variants not found in the opposite S6_MMP-3_ sort are colored in gray, and the ones found in both sorts are colored by the MMP-3 ER of the opposite sort. Regions where high-affinity specific variants are located are lower right for S6_MMP-3 HIGH_ and upper right for S6_MMP-3 LOW_. Squares indicate 5 variants selected based on the S6_MMP-3,HIGH_ sort, and triangles -4 variants based on the S6_MMP-3,LOW_ sort. We then narrowed down the variants to 7 by eliminating variants with similar sequences. D, The sequence logos from the S6 _MMP14_ sort (48) of the two dominating clusters, each including about 35% of all sequences. G66 is conserved in both main clusters and many other smaller clusters. Cluster 1 shows further conservation of R67_D_ while cluster 2 shows conservation of G67_B_/G67_C_. Note that positions 67 and 68 are conserved in both clusters but the library design allows only two amino acids at these positions.

We next sorted the N-TIMP2 library for high expression and high MMP-14 binding using diagonal gates collecting the top 1-2% binding/expression variants at decreasing MMP-14 concentrations for seven rounds. (Supplementary Fig. S3). After significantly enriching binders to MMP-14 (Fig. 2B, left), we searched for N-TIMP2 variants that bind with high affinity to our target MMP-14 but bind weakly to our off-target, MMP-3 by utilizing NGS. For this purpose, the clones from sort S5 were sorted to collect high-affinity binders to MMP-14 (sort S6_MMP-14_; Fig. 2B, center). In a parallel experiment, we collected N-TIMP2 variants that bind with high affinity or low affinity to MMP-3 in sorts S6_MMP-3,HIGH_ and S6_MMP-3,LOW_, respectively (Fig. 2B, right).

We then performed NGS on the four sorted populations (S5, S6_MMP-14_, S6_MMP-3,HIGH,_ and S6_MMP-3,LOW_) in duplicates obtaining 10^6^ reads per sample. Enrichment ratios (ER) were calculated as the ratio of variant frequency appearing in the S6 population (S6_MMP-14_, S6_MMP-3,HIGH_ or S6_MMP-3,LOW_) and its frequency in the S5 population. The errors in variant frequency and ER were estimated by first calculating the error vs average count relationship in representative independent biological duplicate experiments and then using the same relationship for experiments for which no biological duplicates were available (See Methods and Supplementary Fig. S4). We analyzed ERs for all N-TIMP2 variants present in the S6 sorts and looked for variants that were simultaneously enriched in the S6_MMP-14_ sort (ER>1) and the S6_MMP-3,LOW_ sort (highest ER) or were enriched in the S6_MMP-14_ sort (ER>1) sort but were depleted in the S6_MMP-3,HIGH_ sort (lowest ER) (Fig. 2C). We narrowed down our selection to seven such variants representing diverse loop sequences and exhibiting low errors in ERs (Table 2). These variants were analyzed and clustered (Fig. 2D), showing two populous clusters dominating the sequence landscape, which have distinctive motives: cluster 1 had a conserved R at position 67_D_, while cluster 2 had a conserved G and D at positions 67_B_ and 67_C_, respectively.

**Table 2.** Sequences of all variants selected by NGS.

|  | 66 | 67 | 67<sub>A</sub> | 67<sub>B</sub> | 67<sub>C</sub> | 67<sub>D</sub> | 67<sub>E</sub> | 68 | 71 |
| --- | --- | --- | --- | --- | --- | --- | --- | --- | --- |
| WT | A | P |  |  |  |  |  | S | V |
| Var1 | G | A | K | G | D | A | A | G | I |
| Var2 | G | A | K | G | D | P | E | G | A |
| Var6 | G | D | T | E | A | R | M | G | M |
| Var3 | G | A | K | D | D | P | A | G | M |
| Var4 | G | D | T | S | S | S | P | G | M |
| Var5 | G | A | D | D | T | P | S | G | Q |
| Var7 | S | D | K | E | G | R | P | G | G |

### Variant characterization

The seven selected variants were individually expressed on yeast and the apparent binding affinities (K_D_s) of these variants for MMP-14 and MMP-3 were determined using YSD titrations (Fig. 3A). Figure 3A shows that all the selected variants exhibited slightly improved K_D_ toward MMP-14 relative to WT N-TIMP2 as expected. At the same time, all the selected N-TIMP2 variants exhibited considerably worse K_D_ to MMP-3 in comparison to WT N-TIMP2, with some variants showing no binding to MMP-3 at concentrations tested (up to 1 µM). Thus, in YSD titration experiments, all seven variants showed enhanced binding specificity toward MMP-14 vs. MMP-3 as desired. Furthermore, the apparent affinities from the YSD titrations correlated with the ERs calculated by NGS (Supplementary Fig. S5).

**Figure 3.**
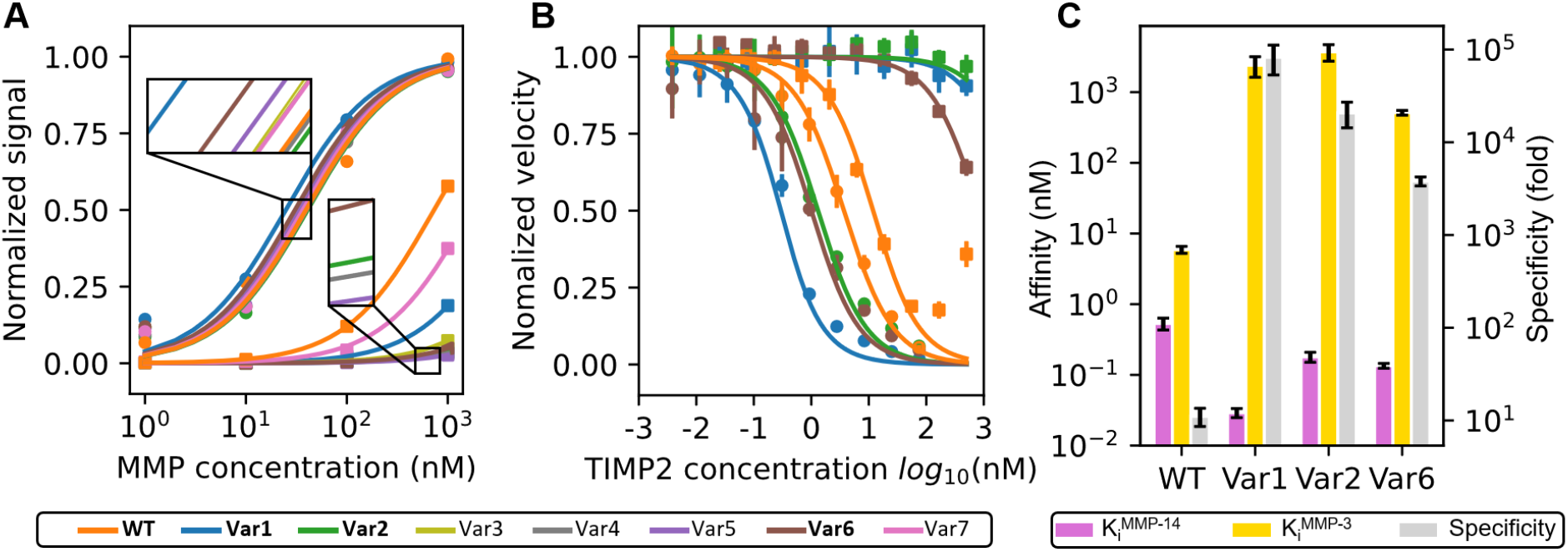
Evaluation of affinity and specificity of the selected N-TIMP2 variants. A, YSD titration of the 7 NGS-selected specific variants. MMP-14 titration data points are shown in circles and MMP-3 data points in squares. B, MMP inhibition assay for the three purified specific N-TIMP2 variants. The experiment was done in triplicates and each replicate was fitted individually. Error bars are standard deviations. C, variant K_i_ values are plotted for MMP-14 and MMP-3. Specificity is calculated as 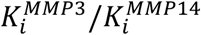. Error bars are standard deviations.

Three N-TIMP2 variants that showed high specificity increase in YSD experiments were chosen for further characterization: the highest affinity variant for MMP-14, Var1, and two variants that showed low affinity to MMP-3, Var2 and Var6 (Table 2).

These three variants as well as WT N-TIMP2 were expressed in Pichia pastoris and purified as previously described (30). To determine the inhibition constant (K_i_) of these N-TIMP2 variants for MMP-14 and MMP-3 (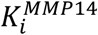 and, 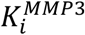 respectively) we performed MMP activity assays that measure cleavage of an MMP fluorogenic substrate as a function of inhibitor concentration (49) (Fig. 3B). Our data shows that WT N-TIMP2 exhibits K_i_ values of 0.53 ± 0.1 nM and 5.9 ± 0.7 nM for MMP-14 and MMP-3, respectively, corresponding to 11-fold specificity (i. e. 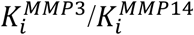) toward the target MMP-14 relative to the off-target MMP-3. The three N-TIMP2 variants showed 3-18 times improvement in 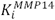, and 86-630 times worse 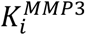 compared to WT N-TIMP2 (Fig. 3C, Table 3). Our best variant, Var1 showed 83000-fold specificity toward MMP-14 relative to MMP-3, 7500-fold more specific than WT-TIMP2. In addition to testing binding specificity for the two MMPs directly used in our experiment as positive and negative targets, we measured their K_i_ to another off-target MMP, MMP-9 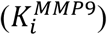. Our data show that variants are 15-66 fold more specific than WT-TIMP2 toward MMP-14 relative to MMP-9, even though MMP-9 was not used as off-target in the engineering process.

**Table 3.**
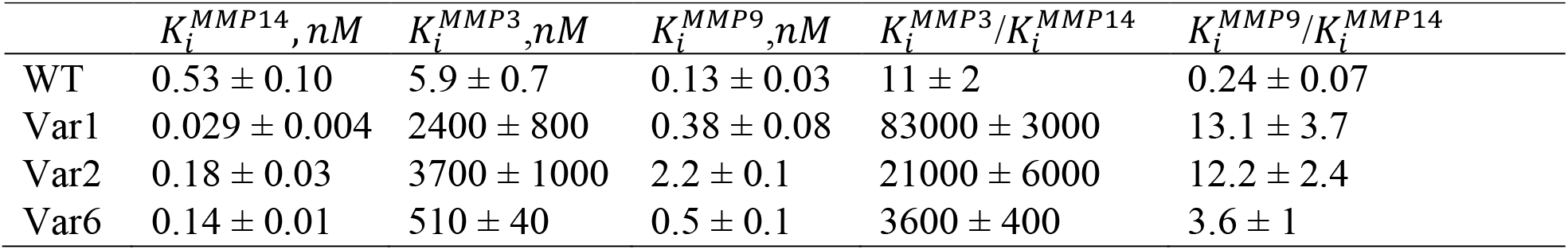
MMP-14, MMP-3, and MMP-9 inhibition constants of the engineered variants.

### Modeling of N-TIMP2 variant interactions with MMP-14 and MMP-3

To better understand the reasons for the increased binding specificity of the selected variants toward MMP-14 relative to MMP-3, we attempted to build structural models of these N-TIMP2 variants interacting with MMP-14 using two different methods, AlphaFold (AF) multimers (50) and Rosetta FKIC(45). We first focused on Var1, our most specific variant. While most of the Var1/MMP-14 complex structure was modeled with high accuracy with AF, the most interesting part of the molecule, the engineered loop region, was modeled with low confidence (pLDDT score of about 40). This was not surprising since AF was trained to infer interacting residues from a multiple sequence alignment (MSA)(51) and the engineered loop is dissimilar from sequences in the AF training set. However, we possessed additional information, the NGS dataset for loop sequences that bound MMP-14 and we thought that this data could help us overcome modeling difficulty. As all loop sequences in the same cluster are likely to share the same conformation, we decided to include our NGS data for loop sequences in cluster 2 (Fig. 2D), to which Var1 belongs, into the MSA alignment, producing the AF custom MSA model (AF-NGS). This model resulted in a dramatic increase in the predicted accuracy of the engineered loop (pLDDT score of 85), producing the same accuracy as observed for similar length loops in the modeled complex (Supplementary Fig. S6A). In a second approach, we used the Rosetta FKIC protocol to build the engineered CD loops into the WT N-TIMP2/MMP-14 complex (see Methods). This approach showed two alternative low-energy loop conformations, named RL1 (the top scoring) and RL2 (the second top scoring) models (Supplementary Fig. S7) with the RL1 model similar to the AF model and extending towards the non-conserved residue G210 on MMP-14, and the RL2 model similar to the AF-NGS model and extending towards the non-conserved residue E248 on MMP-14 (Fig. 4A, Fig. 4B, Supplementary Fig. S8A, Supplementary Fig. S8B). Despite extending in opposite directions, the models have local similarities with perfectly superposing fragments. Remarkably, the fragment composed of D67_C_, A67_D_, and A67_E_ is also found in the 7 designs used for the design of the library (Supplementary. Fig. S8C).

**Figure 4.**
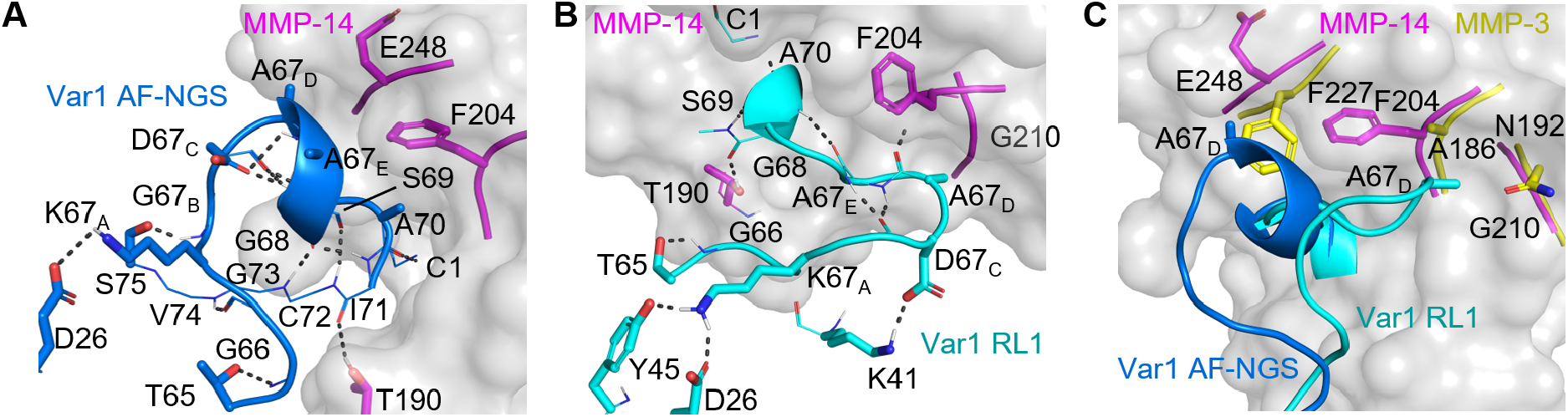
Modeling N-TIMP2-Var1 loop interactions with MMP-14 and MMP-3. MMP surface is shown in light gray, MMP-14 interacting residues are shown in magenta and MMP-3 interacting residues shown in yellow, while the engineered N-TIMP2 loop is shown in blue according to AF-NGS model and in cyan according to RL1. A, Var1 interactions with MMP-14 according to AF-NGS, showing intermolecular hydrophobic interaction of A67_E_/A67_D_ with F204 on MMP14, and intermolecular I71-T190 H-bond. 13 intramolecular H-bonds and the K67_A_-D26 salt bridge stabilize the loop conformation, showing no cavities both at the binding interface and in the loop; B, Var1/MMP14 complex according to the RL1 model showing the intermolecular hydrophobic interactions of A67_E_/A67_D_ with F204 on MMP14 and interface H-bonds A67_D_-F204 and I71-T190; 9 intramolecular H-bonds and the K67_A_-D26 and D67_C_-K41 salt bridges that stabilize the loop conformation. C, AF-NGS model (dark blue) showing incompatible apparent clashes with F227 on MMP-3, and RL1 model (cyan) showing apparent clashes with N192 on MMP-3.

To further evaluate different models, we decided to compute their compatibility with the sequence profile found by NGS. Var1 belongs to cluster 2 of NGS sequences, with the conserved G/D motif at positions 67_B_ and 67_C_. By applying the recent protein design methodology protein-MPNN(52) or Rosetta FastDesign(43) we could recapitulate the sequence profile observed in NGS for the AF-NGS model and the similar RL2 model. The agreement between the conserved positions in the NGS and the designed loop sequences was less while using the RL1 model as an input, and much worse for the AF single loop model, where many amino acids were allowed (Supplementary Fig. S9). This shows that the AF-NGS and RL2 models better explain the NGS data (Fig. 4A). Next, we similarly modeled the interaction between MMP-14 and Var6, whose engineered loop belongs to sequence cluster 1. As with Var1, AF model produced low accuracy for the loop region when only one loop sequence was modeled (pLDDT score of 40); this accuracy increased significantly when AF with custom MSA protocol was applied (pLDDT score of 60-65, Supplementary Fig. S6B). Also similarly, the conservation of R at 67_D_ was recapitulated after designing sequences based on the AF-NGS model, while more amino acid diversity was observed using the AF model, demonstrating the superiority of the AF with custom MSA protocol for loop modeling (Supplementary Fig. S10).

Thus, modeling of variant Var1 showed two alternative loop conformations in the best models: AF-NGS (Fig. 4A) and RL1 (Fig. 4B). Further inspection of the models with MMP-14 showed that in both models the engineered CD loop is tightly packed against MMP-14, making some new favorable hydrophobic interactions between A67_E_/A68_D_ and F204 on MMP-14, and interface hydrogen bonds with T190 on MMP-14. Both loop models are stabilized by the K67_A_-D26 salt bridge and an intricated network of intramolecular H-bonds, particularly in the AF-NGS model, where all residues are involved in one or more H-bonds (Fig. 4 A-B).

Thus, the models suggest that an increase in buried hydrophobic surface area, interface H-bonds, and high loop stabilization are likely responsible for the increased affinity of Var1 to MMP-14. To model Var1 in apparent complex with MMP-3, we superposed Var 1 from the Var1/MMP-14 structural models on the structure of the WT N-TIMP2/MMP-3 complex modeled with AF. Our analysis showed that without any change in the loop conformation, variants Var1 CD loop would clash with either F227 (in AF-NGS) or with N192 (in RL1) on MMP-3 (Fig. 4C). These apparent clashes do not occur in the Var1/MMP-14 interaction since smaller residues are present at positions corresponding to F227 and N192 and the backbone conformations exhibit slight differences. Var1 also contacts F204 on MMP-14, which is not present in MMP-3. Modeling of Var1/MMP-3 complex by Rosetta FKIC did not show any loop conformations similar to what is observed in the Var1 interaction with MMP-14, suggesting that substantial side chains repacking and backbone rearrangement in MMP-3 or the Var1 loop would be needed for binding; these rearrangements might result in Var1 considerably worse affinity to MMP-3. Similar conclusions were reached with respect to Var2, whose loop sequence belongs to the same cluster as Var1 (Supplementary Fig. S11A). Var6 belongs to loop cluster 2 and shows a different loop conformation (Supplementary Fig. S11B), which is stabilized by 9 intramolecular hydrogen bonds and the R67_D_-D67 salt bridge. Superposition of a Var6 model with the structure of WT N-TIMP2/MMP-3 complex shows that the loop is interacting mostly through hydrophobic interactions with non-conserved residues on MMPs; different strengths of these interactions with MMP-14 and MMP-3 could be the reason for the observed enhanced specificity of this variant.

### Evaluating loop models through MMP-3 mutagenesis

Our modeling suggested two alternative conformations for the engineered CD loop of N-TIMP2 Var1, our most specific variant. In the AF-NGS model, it is apparently clashing with F227 on MMP-3 while in the RL1 model it is clashing with N192 (Fig. 4C). To establish which of the loop conformations could occur in reality, we designed two MMP-3 mutants that substitute the predicted clashing residues with a glycine thus possibly relieving the clash and rescuing binding of N-TIMP2 Var1 to MMP-3 in only one predicted loop conformation. Glycine was chosen as a substitution at both positions because it is small and is present in other MMP homologs at these positions. MMP-3 F227G would likely rescue binding of Var1 in the AF-NGS modeled loop conformation (Fig. 4A) but would be a neutral mutation in the RL1 predicted conformation (Fig. 4B). On the contrary, MMP-3 N192G would likely rescue binding of Var1 in the RL1 loop conformation but would be neutral in the case of the AF-NGS loop conformation (Fig. 4C). Both single MMP-3 mutants were experimentally constructed and tested for inhibition by N-TIMP2 Var1. Fig. 5 shows that WT MMP-3 is fully inhibited by WT N-TIMP2, but not by Var1 at concentrations of 30 and 100 nM, in agreement with the results shown in Fig. 3A and B. Similarly, MMP-3 N192G is completely inhibited by WT N-TIMP2, but not by Var1. In contrast, MMP-3 F227G shows concentration-dependent restoration of inhibition by Var1. This implies that the apparent clash with F227 is one of the main reasons for low MMP-3 affinity of Var1, and is consistent with the F227G design removing the clash thus rescuing binding. These mutagenesis results strongly suggest that the AF-NGS model shown in Fig. 4A is correct.

**Figure 5.**
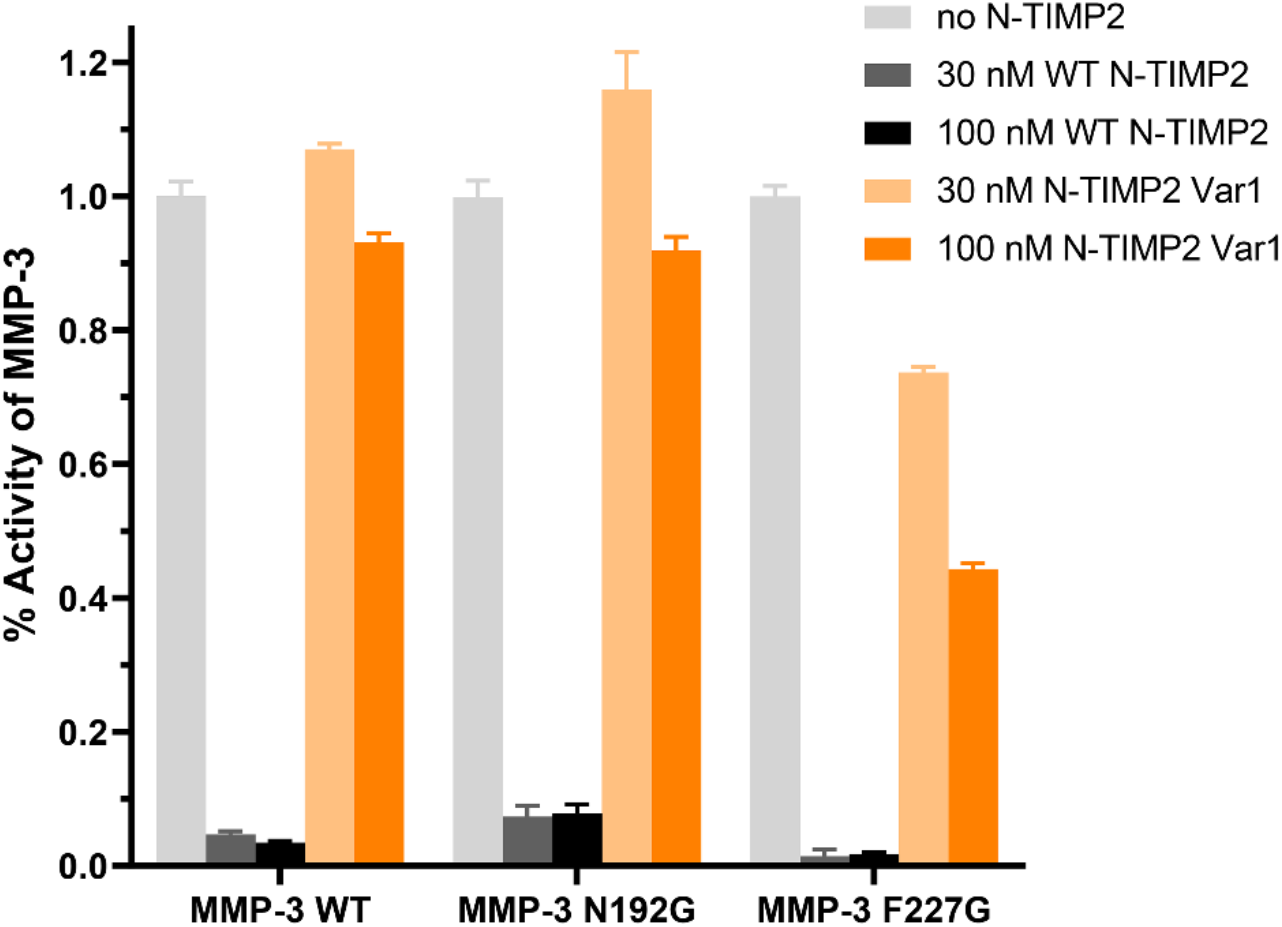
Inhibition of MMP-3 variants by WT N-TIMP2 and Var1. For each MMP-3 variant, fractional residual activity for cleavage of a thiopeptolide substrate is shown following 1 h incubation with the indicated N-TIMP2 WT or variant concentrations, in comparison with the activity for the uninhibited enzyme. MMP-3 WT and N192G show little susceptibility to inhibition by 30 or 100 nM N-TIMP2 Var1, whereas MMP-3 F227G shows concentration-dependent restoration of inhibition by N-TIMP2 Var1. Error bars represent SD for triplicate measurements.

### Testing N-TIMP2 Var1 inhibition of breast cancer cells invasion

We next measured the ability of WT N-TIMP2 and Var1 to inhibit the invasion of breast cancer cells in Matrigel transwell invasion assays. For this purpose, we used two different cell lines, BT549 and MDA-MB-231, both of which are common models of triple-negative breast cancer and are known to overexpress MMP-14. In BT549 cells, treatment with 500 or 1000 nM of WT N-TIMP2 or the N-TIMP2 variant Var1 resulted in significant reduction of cellular invasion (Fig. 6A). Var1 showed significantly enhanced ability to inhibit invasion relative to WT N-TIMP2 (p=0.0375), as assessed by two-way ANOVA. Experiments with MDA-MB-231 cells also showed a trend of reduced invasion when cells were treated with WT N-TIMP2 or Var1, although the magnitude of the effect was more modest (Fig. 6B). Here, significant inhibitory effects were found only in the condition treated with 1000 nM of Var1. We conclude that Var1 fully retains the capability of WT N-TIMP2 to inhibit the invasion of triple-negative breast cancer cells, in some cases with significantly increased potency relative to the WT N-TIMP2. The effects observed with the different cell models may reflect differences in the spectrum of MMP activities that promote invasion of different tumors, since although MMP-14 is a known critical mediator of invasion, additional MMPs including MMP-9 can also play a role in invasion of triple-negative breast cancer (53).

**Figure 6.**
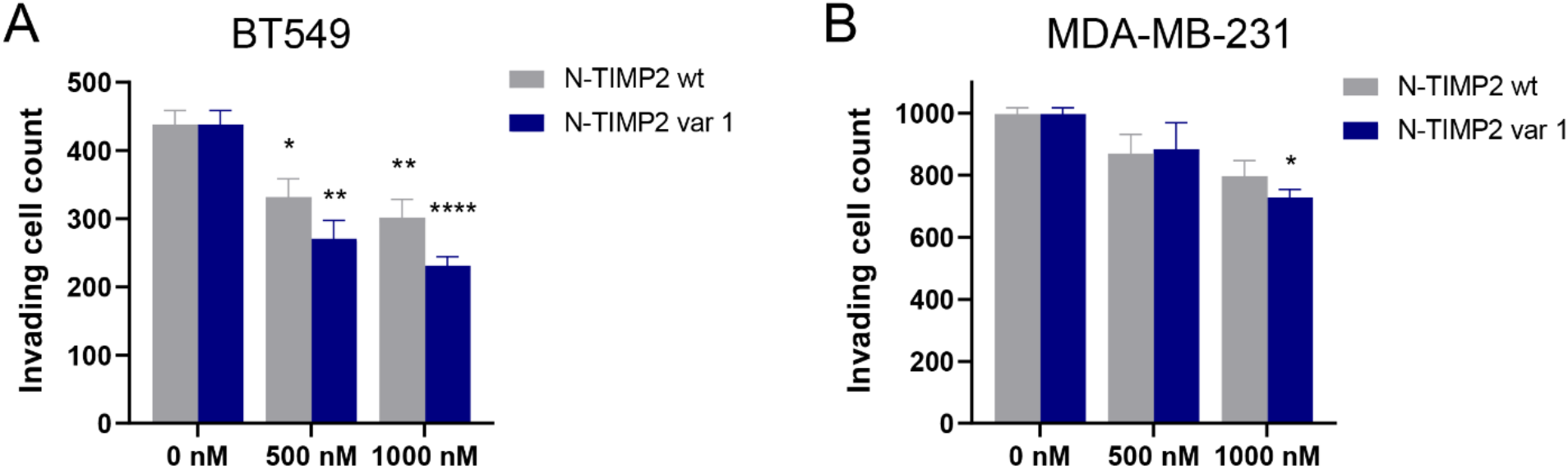
Inhibition of triple-negative breast cancer cell invasion by N-TIMP2 variants. A, in Matrigel transwell invasion assays, treatment of BT549 cells with 500 or 1000 nM of WT or variant N-TIMP2 significantly reduced invasion relative to vehicle-only treated controls (0 nM). Greater inhibitory effects were seen for the N-TIMP2 variant compared to WT N-TIMP2 (p=0.0375, 2-way ANOVA). B, in Matrigel transwell invasion assays, treatment of MDA-MB-231 cells with WT or variant N-TIMP2 shows a trend of reduced invasion relative to vehicle-only treated controls (0 nM), and a significant reduction found for treatment with 1000 nM of the N-TIMP2 variant. Graphs show mean and SEM for quadruplicate biological replicates. *, p<0.05; **, p<0.01; ****, p<0.0001 (multiplicity-adjusted p-values with Dunnett correction for comparison to the untreated control condition).

## Discussion

In this study, we developed a new strategy for engineered specificity into non-specific MMP inhibitor N-TIMP2 by extending the CD loop of TIMP2 to introduce interaction with the non-conserved region of MMP-14. All the previous attempts to engineer TIMPs mutated some interfacial TIMP positions, however, they did not explore insertions or deletions in the TIMP sequence. Such modifications, however, are frequently used by natural proteins and/or antibodies to acquire high affinity and specificity for their targets (54). Such a strategy has also been used when designing orthogonal colicin/immunity pairs with high specificity for each other (40). TIMPs have loops comparable in structure to an antibody (55). Thus, we postulated that applying a strategy used in antibody affinity maturation would also work in TIMP engineering. Indeed, our results show that specificity increase achieved through TIMP loop extension by far exceeds the increase achieved in previous TIMP engineering studies that introduced a number of disjoint mutations in the TIMP binding interface.

Our best variant N-TIMP2 variant, Var1, with 5-residue insertion and 4 additional mutations, bound MMP-14 with 83000 times better affinity compared to off-target MMP-3. This huge affinity difference is sufficient to provide selectivity toward MMP-14 *in vivo*. The increase in binding specificity came from both a medium-size improvement in affinity to MMP-14 (18-fold) and a large 410-fold reduction in affinity to the off-target MMP-3, resulting in 7500-fold improvement in binding specificity. In comparison, an N-TIMP2 mutant with disjointed mutations at the binding interface, previously engineered by our group and collaborators, exhibited 900-fold improved affinity for MMP-14 but also showed improved affinity to four off-target MMPs, thus demonstrating moderate enhancement in binding specificity (31). Another N-TIMP2 variant that was engineered in a competitive YSD setup with positive and negative target MMPs showed 57-fold increase in specificity for MMP-14 relative to an off-target MMP-9 and only ∼3-fold improvement in binding affinity (30).

In similar approaches, N-TIMP2 has been engineered for enhanced specificity toward other MMP targets such as MMP-1 (28), MMP-3 (56), and MMP-9 (30) by randomizing several N-TIMP2 positions and using competitive directed evolution approaches. In all these studies, the introduction of several mutations resulted in increased specificity but slightly decreased binding affinity to the target MMP. Thus, our approach of extending one of N-TIMP2’s loops presents a more attractive strategy for enhancing both N-TIMP2 affinity to the target MMP and enhancing its binding specificity. Moreover, the new loop extension on N-TIMP2 could be combined with previously identified single mutations that enhance binding specificity towards MMP-14, most of which are far from the engineered loop (28, 30, 31).

Our N-TIMP2 variants exhibit higher affinity to MMP-14 (K_i_ of 0.03 nM) compared to specific antibodies that have been developed for MMP-14 (K_i_ of 0.8 nM (17), 9.7 nM (18), and 45 nM (20)). Some of these antibodies showed promising preclinical results by decreasing tumor progression in mice breast cancer xenografts (17). Most of these antibodies bind to exosites or in the proximity of the active site but without directly inhibiting the catalytic zinc (14). Our N-TIMP2 variants have several advantages over antibodies. Specifically, they bind to the active site of MMPs and hence achieve complete MMP inhibition at saturating concentrations unlike some of the developed anti-MMP antibodies (15). In addition, their relatively small size allows them to penetrate tissues better than antibodies and they might be less immunogenic compared to antibodies. Furthermore, full-length WT TIMP2 has already shown promise in preclinical studies in a murine model of triple-negative breast cancer (27).

To select MMP-14 specific variants, we used two sorts for our negative target MMP-3, S6_MMP-3,HIGH_ and S6_MMP-3,LOW_, with the goal of selecting mutants that are depleted in the first sort and enriched in the second sort (Fig. 2C). Interestingly, quite a few variants are found in one sort but not in the other. Of importance, variants not found in the S6_MMP-3,HIGH_ sort are among the most specific in the S6_MMP-3,LOW_ sort (gray points located in the upper part of the right plot on Fig. 2C, representing relatively specific variants). However, the opposite is not true as variants not found in the S6_MMP-3,LOW_ sort are not the most specific in the S6_MMP-3,HIGH_ sort (gray points located on the mid-upper part of the left plot of Figure 2C, representing relatively non-specific variants). One of the reasons for this fact is that the S6_MMP-3,LOW_ sort is enriched for high-specificity variants, while the S6_MMP-3,HIGH_ sort is depleted of specific variants resulting in a lower count. Application of a threshold during the NGS data analysis removes variants with low counts, which are among the specific ones in the S6_MMP-3,HIGH_ sort. The S6_MMP-3,LOW_ sort appears to discriminate low MMP-3 binding better than the S6_MMP-3,HIGH_ sort, with variants selected from the S6_MMP-3,LOW_ sort showing the lowest binding to MMP-3 (Supplementary Fig. S5 and Fig. 3A). Yet, multiple variants that were depleted in the S6_MMP-3,HIGH_ sort were identified (Fig. 2C left); among these was our best variant Var1 (Supplementary Fig. S5). The S6_MMP-3,HIGH_ and S6_MMP-3,LOW_ sorts therefore provided complementary information that could also be combined for the validation of our loop engineering protocol.

To explain the reason of variant’s enhanced binding specificity toward MMP-14 relative to MMP-3, we constructed models of the engineered loop/MMP interactions using different modeling approaches. Modeling of the designed loop proved to be difficult with standard AF methodology since the loop has no sequence homology to any protein in the PDB. Hence, we designed new methodology that incorporated the NGS data on selected loop sequences compatible with high-affinity binding to construct a custom MSA for the AF input. Using this approach, we were able to predict the conformation of Var1 loop with high accuracy and to further validate this model computationally and with mutational analysis. Comparison of the loop models in characterized variants to the modeled structures of the initial designs (Fig, 1D) demonstrated the advantages of focusing the loop library through the design process. For example, positions 67_D_ and 67_E_ were shown to contact the non-conserved MMP surface in the 7 loop designs and in the variants found experimentally (Fig. 1D, Supplementary Fig. S8C); this finding justifies full randomization of these positions in our library. The loop end positions 67 and 68 were shown to be spatially restricted and hence were focused to only two amino acid choices. And the insertion points at the beginning and the end of the loop (T65, S69, and A70), maintained the structure observed in WT-TIMP2 (Fig. 1B) in the models of NGS-selected variants (Fig. 4A, Fig. 4B), supporting the choice of the insertion location. Furthermore, while the initial models of the designed loop prior to experiments differ from the AF-NGS model of the experimentally selected highly specific loops, the two models exhibit local similarities, for example in the D67_C_, A67_D_, and A67_E_ fragment, thus justifying the library focusing based on computational designs.

Our modeling suggests that an increase in hydrophobic surface area and introduction of intra- and inter-molecular H bonds results in enhancement in binding affinity to MMP-14 as such interactions are correlated with binding affinity (58). The reduction of affinity to MMP-3 comes from an apparent clash between the loop and F227 on the enzyme and no similar loop conformations are allowed in the complex with MMP-3 (Fig. 4C). Removal of this clash could restore high affinity of Var1 to MMP-3 as was confirmed with mutagenesis experiments (Fig. 5). The introduction of apparent clashes with off-targets is an attractive strategy for specificity design, which has been used in a number of previous studies (59, 60). Besides contacting non-conserved regions, N-TIMP2 Var1 loop model appears to have an intricate high-density network of intramolecular hydrogen bonds (Fig. 4A) which are unlikely to occur in other loop conformations that allow binding to MMP-3, contributing to Var1 specificity.

In conclusion, using computational design, YSD and NGS, we developed an N-TIMP2 variant with a CD-loop extension that inhibits MMP-14 with exceptionally high affinity and specificity. This variant exhibited improved compared to WT N-TIMP2 inhibition of cell invasion in cancer cell lines that overexpress MMP-14. The general approach of N-TIMP2 loop extension and introduction of interactions with non-conserved MMP sites is a general methodology that could be applied to engineer N-TIMP2 variants targeting other MMP family members.

## Methods

### Design of the loop extension

We built 2000 loop models with Rosetta Remodel, with 7 full random positions (67-73 Rosetta numbering). The flanking positions were flexible but not designable. At this design phase, we also did not allow repacking outside of the loop, to find models with high complementarity to the natural most stable conformation of MMP-14. We hierarchically clustered the models with a 1 Å RMSD distance threshold (measured over the loop backbone, with superposition of the rest of the N-TIMP2), using the Python library scipy. This resulted in 26 clusters, which were visually inspected and the clusters in contact with target residues on MMP-14 were analyzed. One of these clusters was selected to design single designs. First, we expanded the cluster by building ten thousand additional loop models with Cα atoms constrained to be within 2 Å from a representative model close to the cluster center. All designable residues Cα were restrained to corresponding fixed atoms. The resulting models were analyzed for the per-residue binding energies of the target residues that the original cluster was targeting, in this case, P207 and G210. The per residue binding energy was calculated as the per residue energy in the bound conformation minus the per residue energy unbound conformation. Therefore, the best 35 designs were selected based on G210 and P207 per residue energies and the loop model total energy. For each of the 35 selected designs, we run Rosetta FastDesign as a mover. Only designed loop residues (67-73) and flanking positions were allowed backbone movements. Neighbor amino acids within 8 Å from any designed position were allowed to repack, both on N-TIMP2 and on MMP-14. Designable amino acids were restricted by disallow_nonnative_loop_sequences, and FastDesign mover was run with the relaxscript “InterfaceDesign2019”. 30 independent trajectories were run for each of the 35 designs. By inspecting the resulting models, we noticed that in some models Y36 in MMP-14 was not coordinating a structural calcium ion like in the original structure, and the energy decreased after reversion. So, we set these positions to their original sidechain conformations in the crystal structure 1BUV and disallowed repacking in subsequent steps. We ran the Rosetta kinematic loop closure with fragment (FKIC) (44) protocol for the 23 unique loop sequences (from the selected 35 designs) both in the MMP-14 bound and unbound conformations. We ran Rosetta protocols on the HPC of the Hebrew University using the slurm batch scheduler.

### WT N-TIMP2/MMP14 and WT N-TIMP2/MMP-3 models

Modeling of N-TIMP2 variant interactions with MMP-14 was based on the crystal structure for the WT N-TIMP2/MMP-14 complex (PDB ID: 1BUV), which is bovine TIMP2 and human MMP14. TIMP-2 was humanized by Rosetta FastDesign by taking the best of 50 runs, or AlphaFold multimers (50) since it became available. The N-TIMP2/MMP-3 complex was modeled either by aligning the unbound MMP-3 structure (PDB ID: 6MAV (61)) to the highly homologous TIMP2/MMP10 structure (PDB ID: 4ILW (62)) or using AlphaFold multimer. These two N-TIMP2/MMP-3 models were generally similar but exhibited a slightly different N-TIMP2 binding orientation relative to MMP-3 (Supplementary Fig. S13).

### Modeling of Var1 Var2 and Var6 in complex with MMP-14 and MMP-3

We modeled variant Var1 in the N-TIMP2/MMP-14 complex using Rosetta kinematic closure with fragments (FKIC) and AlphaFold multimers while Var6 was modeled with AlphaFold multimers only and Var2 with FKIC only. For FKIC fragments were picked with Robetta Server, and we run 5000 trajectories per variant, hierarchically clustered the models based on the loop RMSD cutoff of 2-4 Å, and inspected the lowest energy model from the top5 clusters with lowest energy models (the five bigger size marks in Supplementary Fig. S7). Besides Var1 and Var2, another 4 close neighbors were modeled both in the MMP-14 and MMP-3 complexes (loop seq: GAKGDAAGSAM, GAKGDPAGSAV, GAVGDAAGSAI, GAKGDPEGSAV).

The RL1 and RL2 conformations were both found in all neighbors as top5 clusters, and consistently ranked as the top models. For the modeling with AlphaFold multimers, we used the colab implementation (63), uploading a custom MSA, which contained the additional sequences from the NGS data (both single and paired). We used sequences present in both S5 and S6_MMP-14_ with count threshold of 100. We tried different sequence selections based on clustering with BLOSUM62 distance, minimum ER for MMP-14, and minimum BLOSUM62 distance from the modeled sequence. The minimum BLOSUM62 distance was used to select sequences more likely to have the same conformation as the model sequence, and the minimum ER for MMP-14 to discard sequences that might have very low binding. For Var1 we obtained the best accuracy with a minimum BLOSUM62 distance of 28 (over all 11 loop amino acids) and ER MMP14 > 0.14, which resulted in about 100 sequences. For Var6 we use sequences in cluster 1 (BLOSUM62 distance threshold 40) restricted by a minimum BLOSUM62 distance of 28 which resulted in about 100 sequences, and we did not optimize further.

### Evaluation of models through sequence design protocols

We use protein-MPNN(52) and Rosetta FastDesign to evaluate the models by their ability to recapitulate the NGS data. We hypothesized that the amino acids that are conserved in the NGS clusters must have a core role that is independent of the identity of the variating ones. For protein-MPNN, we analyzed the returned conditional probability of conserved amino acids in the loop, given the sequence only outside of the loop and the model structure. For Rosetta FastDesign, we perform single-position mutational scans giving the sequence and structure of all other positions, running 20 trajectories per position, and then taking the average score of the best 5.

### Library construction and yeast transformation

To transform the library we used the plasmid gap repair by homologous recombination strategy (64), in which a linearized vector and a library insert have homologous ends, and are co-transformed in yeast where they recombine.

The library was synthesized as a mixed-bases-ultramer (Integrated DNA Technologies) harboring all the variation. The ultramer was extended by isothermal primer annealing-extension, using a standard Q5 (New England Biolabs) PCR reaction but holding the temperature at 72 degrees for 30’, which yielded optimal results (Supplementary. Fig. S14A).

To prepare for the yeast transformation, the library was amplified using a standard Q5 PCR reaction but with 2 µM of each primer, which yielded a high quantity of pure product (Supplementary. Fig. S14B). We ran 96 reactions of 100 µl. Reactions were pool-purified with DNA clean-up kit for short fragments (Geneaid) using one column every 10 µg of product, and elution was carried on through 3 columns (about 1µg/µl) and concentrated by ethanol-precipitation.

The pCHA-VRC01-scFv N-terminal yeast display vector (obtained from Dane Wittrup, Massachusetts Institute of Technology) (64) was modified by transfer PCR (65) to insert the N-TIMP2 coding sequence. It was further modified to include a unique NheI cutting site (G|CTAGC) at the site of the engineered CD loop and an appropriate deletion so that once the plasmid is linearized by NheI the ends anneal to the library insert with no overhangs, i.e. the split NheI site sequences (G and CTAGC) are part of the recombined sequence, a property we speculated might increase the transformation efficiency. The vector was linearized with NheI-HF (New England Biolabs) overnight, cleaned up with spin columns, and concentrated by ethanol precipitation. EBY100 was transformed essentially as described (66), using a Gene Pulser II electroporator (Bio-Rad), 3 µg linearized vector, 9 µg library insert, and 350 µl electrocompetent EBY100. This yielded 20-40*10^6^ transformants per electroporation, for a total of 1.9*10^8^ transformants in 6 electroporations. Sequencing of individual variants showed that the loop was incorporated into the gene without errors (14 out of 14 sequenced colonies were correct), and bulk Sanger sequencing of pooled variants showed that all positions had the expected variability.

### Flow Cytometry Analysis and Sorting

FACS analysis and sorting were carried out essentially as described (67), with buffers appropriate for MMP-14 (31) and with minor modifications. Induced EBY100 was concurrently labeled with DyLight-488-MMP-14 (see below) and 1:100 PE Anti-Myc tag antibody (abcam) for 1 h 30’ at room temperature in TNCA (50 mM Tris, pH 7.5, 100 mM NaCl, 5 mM CaCl2, and 1% bovine serum albumin). In any procedure involving sampling of the library (e.g. growing, labeling, sorting, and freezing), we ensured that the 10-100x the input library variability was present. For analysis and YSD titration, 2*106 cells were labeled. To avoid depletion, the labeling volume was such that the molarity of the labeled protein was 10x or more than that of the displayed protein. To achieve that, low concentration labeling was carried out in 10-50 ml volumes. After washing, labeled EBY100 was analyzed or sorted in a FACSAria III (BD Biosciences). For the first library sort (S1) sorting was done at 20 k events/s and sorting mode set to “yield” to scan a total of 4*108 events (2x the initial library variation) in two runs, and the sorts were later combined taking care to maintain the proportion of the collected events in each sort. Subsequent sorts were carried out at 1-10 k events/s and collecting 10-100x the maximum theoretical input library variation (i.e. the number of previously sorted events) and sorting mode set to “purity”. For sorts to be sequenced by NGS, we set the sorting mode to “single-cell” to obtain maximum sort specificity. For affinity maturation sorting we set diagonal sorting gates to collect the top 1-2% binding/expression events for each sort. The MMP-14 concentration was the following: S1 2 µM, S2 2 µM, S3 500 nM, S4 100 nM, S5 20 nM, S6 4 nM. For sorts to be sequenced by NGS, for high binding sorts, we set a diagonal sorting gate at 50% of the maximum binding/expression signal in the library, which resulted in about top 5-10% binding/expression events; for low binding sorts, the gate was set based on a negative control to collect non-binding variants, which resulted in about bottom 4% binding/expression events. S4 was labeled with 20 nM labeled MMP-14 and sorted using a broad gate for high MMP-14 binding/expression to yield S5_NGS_. S5_NGS_ was labeled with 20 nM labeled MMP-14 and sorted for high MMP-14 binding/expression (S6_MMP-14_), or labeled with 1 µM labeled MMP-3 and sorted using a narrow gate for high MMP-3 binding/expression (S6_MMP-3, HIGH_) or sorted for low MMP-3 binding/expression (S6_MMP-3, LOW_).

### NGS: Sample Preparation

Samples for NGS were prepared essentially as described (68)(69). Samples were prepared in duplicates including independent FACS sortings. Plasmids were extracted from 10^8^ cells using Zymoprep Yeast Plasmid Miniprep II (Zymo Research). Samples were treated with ExoI exonuclease (NEB) and lambda exonuclease (NEB) in ExoI buffer to degrade genomic DNA. Samples were cleaned up. The final plasmid number was 30-70*10^6^, determined by the transformants number vs a standard curve in E. coli. All the recovered plasmid was used in a standard Q5 PCR reaction with 18 cycles to amplify and add the Illumina sequencing likers (CS1/2), the product was gel-extracted, and sent for sequencing with an Illumina Miseq in a 2×250 paired-end run obtaining 10^6^ reads per sample, in duplicates.

### NGS: Sequence Analysis

Paired-reads were merged with NGmerge (70), which merges the reads and provides a combined Q_base_ quality score based on an empirical model. All the subsequent analysis was done with custom python scripts. Reads were filtered based on the length to remove shared-run contaminants. The loop sequence was extracted by matching the 5’ and 3’ regions up to 2 edits, and the loop length, and filtered to a minimum of 30 combined Q_base_ quality for all bases.

Biological duplicates were used to calculate variant frequency standard deviations, derived from independent FACS sorts and independent NGS runs. Because the sample differ slightly in the count number, frequencies were used and then transformed back to counts. The centered moving average with a window of 30 of the duplicate frequency standard deviations along the duplicate frequency was calculated, which appeared increasing in absolute terms as previously reported for a more complex statistical model (71, 72). The increasing points of the frequency standard deviation vs frequency curve were transformed to variant count and used to calculate the count standard deviation given the variant count for all other experiments, and the standard deviations propagated in ERs. For ER analysis, a threshold count of 100 was used. Sequences were translated, and variant frequency f_i_ in sorts S_j_ was calculated. Variant frequencies in subsequent sorts were used to calculate enrichment ratios as f_i_ (S_j+1_)/ f_i_ (S_j_). The portion of the ER vs ER plots for high specificity variants was selected by drawing a region of interest, for MMP-14 including variants with ER of about 1 or higher, and for MMP-3 the lowest binding 10-20 variants in either the S6_NGS,MMP-3, LOW,_ or the S6_NGS, MMP-3, HIGH_ sort. Then variants that had worse ER than others both for binding to MMP-14 and not binding to MMP-3 were excluded. From the remaining 9-variants set, we selected 7 variants with divergent sequences to test by YSD titration. Sequences were hierarchically clustered with BLOSUM62 loop distance threshold of 40.

### Yeast titration data analysis

Single variants were expressed on the surface of yeast and YSD titration was carried out essentially as described (64). Median fluorescence was calculated in FlowJo (BD Life Sciences) for PE+ (expressing) cells. Data were fitted in MATLAB Curve Fitting Toolbox (MathWorks) to the rectangular hyperbolic equation for binding:

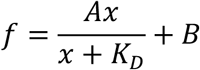

where f is the measured signal and x is the label MMP concentration. A, B, and K_D_ were fitted. The fitting was repeated for each independent replicate and K_D_ values were then averaged. Curves and data were plotted with the python module matplotlib.

### MMP inhibition Assay

MMP inhibition assays for K_i_ studies (Figure 3A and Table 3) were carried out essentially as described (31). Serial dilutions of inhibitor variants were incubated with 0.2 nM MMP-14 or 4 nM MMP-3 for 1h in TNC + 0.05% Brij, and activity was measured by the cleavage of the fluorogenic MMP Substrate MCA-Lys-Pro-Leu-Gly-Leu-DNP-Dpa-Ala-Arg-NH2 (Sigma-Aldrich). Normalized velocities were fitted in MATLAB Curve Fitting Toolbox (MathWorks) to the quadratic equation for binding:

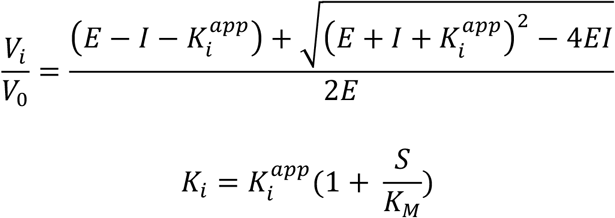

where V_i_ is enzyme velocity at inhibitor concentration I, V_0_ is the enzyme velocity in the absence of inhibitor, E is the active enzyme concentration, S is the substrate concentration, K_M_ is the Michaelis-Menten constant, K_i_ is the inhibition constant, and K_i_^app^ was estimated. The fitting was repeated for each independent replicate and K_i_^app^ values were then averaged. Calculations were performed using a K_M_ value of 1.31 µM for MMP-14, 11.23 µM for MMP-3, and 2 µM for MMP-9 as determined from Michaelis– Menten kinetic experiments performed in triplicate in our laboratory. Curves and data were plotted with the python module matplotlib.

MMP inhibition assays comparing inhibition of WT and mutant MMP-3 (Figure 5) were carried out essentially as described previously (73). MMP molar concentrations were first determined by titration against a reference stock of TIMP-1. Inhibitory activity of N-TIMP2 (WT or variant) toward MMP-3 (WT, F227G, and N192G variants) were determined by measuring reduced rates of hydrolysis of the thiopeptolide substrate Ac-Pro-Leu-Gly-[2-mercapto-4-methyl-pentanoyl]-Leu-Gly-OC_2_H_5_ (Enzo, Life Sciences). Hydrolysis was measured by reaction of the sulfhydryl group with 5,5′-dithiobis(2-nitrobenzoic acid) (DTNB) to form 2-nitro-5-thiobenzoic acid, resulting in an absorbance increase at 412 nm (ε = 13 600/M/cm at pH ≥ 6.0). MMP-3cd (WT, 2 nM; F227G, 0.4 nM; or N192G, 1 nM) was mixed with WT N-TIMP2 or variant (10, 30, or 100 nM) in assay buffer (50 mM HEPES, pH 6.0, 10 mM CaCl_2_, 0.05% Brij-35, 1 mM DTNB) and preincubated at RT for 1 h. Reactions were started by addition of 100 µM thiopeptolide substrate and then followed for 600 s at 410 nm using the Varian Cary UV 100 spectrophotometer (Varian Inc, Palo Alto, CA). Reaction rates were measured in triplicate, fitted by linear regression, and plotted as percentage of residual activity compared to the relevant uninhibited enzyme (WT, F227G, or N192G variant). The complete experiment was repeated twice with consistent results.

### Protein Expression and Purification

The catalytic domain of MMP-14 residues 110-300 was expressed in E. coli KRX cells in inclusion bodies, refolded by dialysis, and purified by a Ni-NTA column and by SEC as previously described (31). The catalytic domain of MMP-3 was produced as described below. Correct protein folding was confirmed by MMP inhibition assays, and active enzyme concentrations were 50-100%.

An expression construct for the proMMP-3 catalytic domain residues 1-255 (74) was mutagenized using primers as follows: MMP-3-F227G 5’gccactccctgggtctcggccactcagccaacactga-3’ and MMP-3-N192G 5’gcccctgggccagggattggcggagatgcccact-3’. Mutagenesis was conducted using the QuikChange multi-site-directed mutagenesis kit (Agilent Technologies, USA) according to the manufacturer’s instructions, and mutated plasmids were confirmed by DNA sequencing (Eurofins, NJ). ProMMP-3 and mutants were expressed in E. coli BL21(DE3), refolded, purified, and activated similarly to our previously described protocol (73). Protein was extracted from inclusion bodies in 8 M urea, 20 mM Tris-HCl, pH 8.6, 20 mM dithiothreitol, and 50 µM ZnCl_2_, and then purified on Q-Sepharose equilibrated with 8 M urea, 20 mM Tris-HCl pH 8.6, and 50 µM ZnCl_2_ and eluted using a linear gradient of NaCl to 0.5 M. Fractions containing proMMP-3 were combined, diluted to A280 of < 0.3mg/ml, and refolded by stepwise dialysis with 50 mM Tris-HCl, pH 7.5, 10 mM CaCl_2_, and 150 mM NaCl. Purified proMMP-3 and mutants were activated for 16 h in the presence of 1 mM 4-amino-phenyl mercuric acetate (Sigma) at 37 °C, and then desalted on PD-10 columns (Cytiva Life Sciences) equilibrated with 50 mM Tris-HCl, pH 7.5, 10 mM CaCl_2_, and 150 mM NaCl. Brij-35 was added to 0.05% and samples were aliquoted and frozen at -80°C until use.

MMPs were labeled with the fluorophore Dylight 488 NHS Ester (ThermoFisher), which labels the proteins at primary amines (-NH_2_), essentially following the manufacturer’s protocol. Dye was dissolved in DMF at 10mg/ml, the labeling buffer was 0.05M sodium borate buffer at pH 8.5, and the reaction was done using 10% dye-DMF, which we speculated was at the limit before DMF could cause protein unfolding. Unreacted dye was removed by ultrafiltration and subsequent dialysis, and the degree of labeling was calculated. The degree of labeling was 2.5 for MMP-14 and 3.7 for MMP-3. The labelling of the proteins was confirmed by SDS-PAGE and imaging under UV.

N-TIMP2 WT and the engineered variants with a C-terminal His-Tag were expressed in the methylotrophic X33 P. pastoris yeast strain using the pPICZα vector (Invitrogen) and purified by Ni-NTA column and SEC as described in our previous work (30).

### Cell Culture and Matrigel Transwell Invasion Assays

Cell cultivation and invasion assays were performed according to procotols that we have previously described (75). BT549 cells were grown in RPMI-1640 media (ATCC) with 10% fetal bovine serum (GeminiBio) and 0.023 U/ml insulin (Sigma). MDA-MB-231 cells were grown in DMEM media (Gibco) with 10% fetal bovine serum (GeminiBio). Cell lines were grown at 37 °C and 5% CO_2_, and the media were changed every two days. Both cell lines were validated by short tandem repeat (STR) genotyping (PowerPlex® 16 HS platform; Promega) of the laboratory seed and distribution stocks.

BioCoat 24-well plate inserts (8.0 micron, Corning Inc.) were coated with 50 µg Matrigel basement membrane matrix in 100 µL of serum-free medium (RPMI 1640 for BT-549 cells; DMEM for MDA-MB-231 cells) and placed at 37 °C for 4 h, and then residual medium was aspirated and replaced with cells (1.5×10^4^ cells per well for BT-549; 5×10^4^ cells per well for MDA-MB-231) suspended in 300 µL of serum-free medium supplemented with 0.1% BSA. The lower invasion chambers contained 750 µL/well of NIH/3T3 cell-conditioned serum-free medium (DMEM containing 50 µg/mL ascorbic acid) as chemo-attractant. Assays were incubated for 18 hours at 37°C in 5% CO_2_. Non-invading cells were then removed from the inserts by scrubbing with a cotton swab, and cells on the lower surface of the filter were fixed with methanol and stained with 0.1% crystal violet. Membranes were dried, photographed at 20x magnification, and counted using the INFINITY ANALYZE software (version 6.5.6, Teledyne Lumenera, Canada).

## Author Contributions

AB conceived the project idea with inputs from JSM. AB performed most of the experimental studies and all the computational studies and wrote the software for the analysis of the NGS data. AB and JMS designed the research. BLW and HA performed the MMP-3 mutagenesis and the inhibition of cell invasion studies. JMS and ESR supervised the project, provided resources, and acquired funding. AB and JMS wrote the manuscript, with contributions from the other authors.

## Acknowledgments

We thank Ora Schueler-Furman and Gideon Schreiber for their insightful discussions. This work was supposed by the US-Israel Binational Science Foundation (BSF) 2017207 (J. M. S.) and NIH R01CA258274 (E.S.R. and J. M.S). In addition, J. M. S. acknowledges the support from ICRF and ISF 3486/20 and the U. of Toronto/HUJI research alliance in protein engineering.

## References

1. Tallant, C., Marrero, A., and Gomis-Rüth, F. X. (2010) Matrix metalloproteinases: Fold and function of their catalytic domains. Biochim. Biophys. Acta - Mol. Cell Res. 1803, 20–28

2. Conlon, G. A., and Murray, G. I. (2019) Recent advances in understanding the roles of matrix metalloproteinases in tumour invasion and metastasis. J. Pathol. 247, 629–640

3. Rydlova, M., Holubec, L., Ludvikova, M., Kalfert, D., Franekova, J., Povysil, C., and Ludvikova, M. (2008) Biological activity and clinical implications of the matrix metalloproteinases. Anticancer Res. 28, 1389–1397

4. Decock, J., Thirkettle, S., Wagstaff, L., and Edwards, D. R. (2011) Matrix metalloproteinases: Protective roles in cancer. J. Cell. Mol. Med. 15, 1254–1265

5. Folgueras, A. R., Pendás, A. M., Sánchez, L. M., and López-Otín, C. (2004) Matrix metalloproteinases in cancer: From new functions to improved inhibition. Int. J. Dev. Biol. 48, 411–424

6. López-Otín, C., and Matrisian, L. M. (2007) Emerging roles of proteases in tumour suppression. Nat. Rev. Cancer. 7, 800–808

7. Sarper, M., Allen, M. D., Gomm, J., Haywood, L., Decock, J., Thirkettle, S., Ustaoglu, A., Sarker, S. J., Marshall, J., Edwards, D. R., and Jones, J. L. (2017) Loss of MMP-8 in ductal carcinoma in situ (DCIS)-associated myoepithelial cells contributes to tumour promotion through altered adhesive and proteolytic function. Breast Cancer Res. 10.1186/s13058-017-0822-9

8. Coussens, L. M., Fingleton, B., and Matrisian, L. M. (2002) Matrix metalloproteinase inhibitors and cancer: Trials and tribulations. Science (80-.). 295, 2387–2392

9. Winer, A., Adams, S., and Mignatti, P. (2018) Matrix metalloproteinase inhibitors in cancer therapy: Turning past failures into future successes. Mol. Cancer Ther. 17, 1147–1155

10. Cathcart, J. M., and Cao, J. (2015) MMP Inhibitors: Past, present and future. Front. Biosci. - Landmark. 20, 1164–1178

11. Lombard, M. A., Wallace, T. L., Kubicek, M. F., Petzold, G. L., Mitchell, M. A., Hendges, S. K., and Wilks, J. W. (1998) Synthetic matrix metalloproteinase inhibitors and tissue inhibitor of metalloproteinase (TIMP)-2, but not TIMP-1, inhibit shedding of tumor necrosis factor-α receptors in a human colon adenocarcinoma (Colo 205) cell line. Cancer Res. 58, 4001–4007

12. Bonadio, A., and Shifman, J. M. (2021) Computational design and experimental optimization of protein binders with prospects for biomedical applications. Protein Eng. Des. Sel. 34, 1–7

13. Fischer, T., and Riedl, R. (2019) Inhibitory Antibodies Designed for Matrix Metalloproteinase Modulation. Molecules. 10.3390/MOLECULES24122265

14. Levin, M., Udi, Y., Solomonov, I., and Sagi, I. (2017) Next generation matrix metalloproteinase inhibitors — Novel strategies bring new prospects. Biochim. Biophys. Acta - Mol. Cell Res. 1864, 1927–1939

15. Appleby, T. C., Greenstein, A. E., Hung, M., Liclican, A., Velasquez, M., Villasenor, A. G., Wang, R., Wong, M. H., Liu, X., Papalia, G. A., Schultz, B. E., Sakowicz, R., Smith, V., and Kwon, H. J. (2017) Biochemical characterization and structure determination of a potent, selective antibody inhibitor of human MMP9. J. Biol. Chem. 292, 6810–6820

16. Shah, M. A., Yanez Ruiz, E. P., Bodoky, G., Starodub, A., Cunningham, D., Yip, D., Wainberg, Z. A., Bendell, J. C., Thai, D., Bhargava, P., and Ajani, J. A. (2019) A phase III, randomized, double-blind, placebo-controlled study to evaluate the efficacy and safety of andecaliximab combined with mFOLFOX6 as first-line treatment in patients with advanced gastric or gastroesophageal junction adenocarcinoma (GAMMA-1). J. Clin. Oncol. 37, 4–4

17. Devy, L., Huang, L., Naa, L., Yanamandra, N., Pieters, H., Frans, N., Chang, E., Tao, Q., Vanhove, M., Lejeune, A., Van Gool, R., Sexton, D. J., Kuang, G., Rank, D., Hogan, S., Pazmany, C., Ma, Y. L., Schoonbroodt, S., Nixon, A. E., Ladner, R. C., Hoet, R., Henderikx, P., Tenhoor, C., Rabbani, S. A., Valentino, M. L., Wood, C. R., and Dransfield, D. T. (2009) Selective inhibition of matrix metalloproteinase-14 blocks tumor growth, invasion, and angiogenesis. Cancer Res. 69, 1517–1526

18. Nam, D. H., Rodriguez, C., Remacle, A. G., Strongin, A. Y., and Ge, X. (2016) Active-site MMP-selective antibody inhibitors discovered from convex paratope synthetic libraries. Proc. Natl. Acad. Sci. U. S. A. 113, 14970–14975

19. Gálvez, B. G., Matías-Román, S., Albar, J. P., Sánchez-Madrid, F., and Arroyo, A. G. (2001) Membrane Type 1-Matrix Metalloproteinase is Activated during Migration of Human Endothelial Cells and Modulates Endothelial Motility and Matrix Remodeling. J. Biol. Chem. 276, 37491–37500

20. Udi, Y., Grossman, M., Solomonov, I., Dym, O., Rozenberg, H., Moreno, V., Cuniasse, P., Dive, V., Arroyo, A. G., and Sagi, I. (2015) Inhibition mechanism of membrane metalloprotease by an exosite-swiveling conformational antibody. Structure. 23, 104–115

21. Sela-Passwell, N., Kikkeri, R., Dym, O., Rozenberg, H., Margalit, R., Arad-Yellin, R., Eisenstein, M., Brenner, O., Shoham, T., Danon, T., Shanzer, A., and Sagi, I. (2012) Antibodies targeting the catalytic zinc complex of activated matrix metalloproteinases show therapeutic potential. Nat. Med. 18, 143–147

22. Brew, K., and Nagase, H. (2010) The tissue inhibitors of metalloproteinases (TIMPs): An ancient family with structural and functional diversity. Biochim. Biophys. Acta - Mol. Cell Res. 1803, 55–71

23. Arumugam, S., Gao, G., Patton, B. L., Semenchenko, V., Brew, K., and Van Doren, S. R. (2003) Increased backbone mobility in β-barrel enhances entropy gain driving binding of N-TIMP-1 to MMP-3. J. Mol. Biol. 327, 719–734

24. Wingfield, P. T., Sax, J. K., Stahl, S. J., Kaufman, J., Palmer, I., Chung, V., Corcoran, M. L., Kleiner, D. E., and Stetler-Stevenson, W. G. (1999) Biophysical and functional characterization of full-length, recombinant human tissue inhibitor of metalloproteinases-2 (TIMP-2) produced in Escherichia coli. Comparison of wild type and amino terminal alanine appended variant with implications for the mechanism of TIMP functions. J. Biol. Chem. 274, 21362–21368

25. Murphy, G., O’Shea, M., Houbrechts, A., Cockett, M. I., Docherty, A. J. P., and Williamson, R. A. (1991) The N-Terminal Domain of Tissue Inhibitor of Metalloproteinases Retains Metalloproteinase Inhibitory Activity. Biochemistry. 30, 8097–8102

26. Huang, W., Suzuki, K., Nagase, H., Arumugam, S., Van Doren, S. R., and Brew, K. (1996) Folding and characterization of the amino-terminal domain of human tissue inhibitor of metalloproteinases-1 (TIMP-1) expressed at high yield in E. coli. FEBS Lett. 384, 155–161

27. Peeney, D., Jensen, S. M., Castro, N. P., Kumar, S., Noonan, S., Handler, C., Kuznetsov, A., Shih, J., Tran, A. D., Salomon, D. S., and Stetler-Stevenson, W. G. (2020) TIMP-2 suppresses tumor growth and metastasis in murine model of triple-negative breast cancer. Carcinogenesis. 41, 313–325

28. Bahudhanapati, H., Zhang, Y., Sidhu, S. S., and Brew, K. (2011) Phage display of tissue inhibitor of metalloproteinases-2 (TIMP-2): Identification of selective inhibitors of collagenase-1 (metalloproteinase 1 (MMP-1)). J. Biol. Chem. 286, 31761–31770

29. Sharabi, O., Shirian, J., Grossman, M., Lebendiker, M., Sagi, I., and Shifman, J. (2014) Affinity- and Specificity-Enhancing Mutations Are Frequent in Multispecific Interactions between TIMP2 and MMPs. PLoS One. 9, e93712

30. Shirian, J., Arkadash, V., Cohen, I., Sapir, T., Radisky, E. S., Papo, N., and Shifman, J. M. (2018) Converting a broad matrix metalloproteinase family inhibitor into a specific inhibitor of MMP-9 and MMP-14. FEBS Lett. 592, 1122–1134

31. Arkadash, V., Yosef, G., Shirian, J., Cohen, I., Horev, Y., Grossman, M., Sagi, I., Radisky, E. S., Shifman, J. M., and Papo, N. (2017) Development of High Affinity and High Specificity Inhibitors of Matrix Metalloproteinase 14 through Computational Design and Directed Evolution. J. Biol. Chem. 292, 3481–3495

32. Arkadash, V., Radisky, E. S., and Papo, N. (2018) Combinatorial engineering of N-TIMP2 variants that selectively inhibit MMP9 and MMP14 function in the cell. Oncotarget. 9, 32036

33. Akiva, E., Itzhaki, Z., and Margalit, H. (2008) Built-in loops allow versatility in domain-domain interactions: Lessons from self-interacting domains. Proc. Natl. Acad. Sci. U. S. A. 105, 13292–13297

34. Rothe, C., Urlinger, S., Löhning, C., Prassler, J., Stark, Y., Jäger, U., Hubner, B., Bardroff, M., Pradel, I., Boss, M., Bittlingmaier, R., Bataa, T., Frisch, C., Brocks, B., Honegger, A., and Urban, M. (2008) The Human Combinatorial Antibody Library HuCAL GOLD Combines Diversification of All Six CDRs According to the Natural Immune System with a Novel Display Method for Efficient Selection of High-Affinity Antibodies. J. Mol. Biol. 376, 1182–1200

35. Schilling, J., Schöppe, J., and Plückthun, A. (2014) From DARPins to LoopDARPins: Novel LoopDARPin design allows the selection of low picomolar binders in a single round of ribosome display. J. Mol. Biol. 426, 691–721

36. MacDonald, J. T., Kabasakal, B. V., Godding, D., Kraatz, S., Henderson, L., Barber, J., Freemont, P. S., and Murray, J. W. (2016) Synthetic beta-solenoid proteins with the fragmentfree computational design of a beta-hairpin extension. Proc. Natl. Acad. Sci. U. S. A. 113, 10346–10351

37. Huang, P.-S., Ban, Y.-E. A., Richter, F., Andre, I., Vernon, R., Schief, W. R., and Baker, D. (2011) RosettaRemodel: A Generalized Framework for Flexible Backbone Protein Design. PLoS One. 6, e24109

38. Hu, X., Wang, H., Ke, H., and Kuhlman, B. (2007) High-resolution design of a protein loop. Proc. Natl. Acad. Sci. U. S. A. 104, 17668–17673

39. Lapidoth, G. D., Baran, D., Pszolla, G. M., Norn, C., Alon, A., Tyka, M. D., and Fleishman, S. J. (2015) AbDesign : An algorithm for combinatorial backbone design guided by natural conformations and sequences. Proteins Struct. Funct. Bioinforma. 83, 1385–1406

40. Netzer, R., Listov, D., Lipsh, R., Dym, O., Albeck, S., Knop, O., Kleanthous, C., and Fleishman, S. J. (2018) Ultrahigh specificity in a network of computationally designed protein-interaction pairs. Nat. Commun. 9, 1–13

41. Ashkenazy, H., Abadi, S., Martz, E., Chay, O., Mayrose, I., Pupko, T., and Ben-Tal, N. (2016) ConSurf 2016: an improved methodology to estimate and visualize evolutionary conservation in macromolecules. Nucleic Acids Res. 10.1093/nar/gkw408

42. Fernandez-Catalan, C., Bode, W., Huber, R., Turk, D., Calvete, J. J., Lichte, A., Tschesche, H., and Maskos, K. (1998) Crystal structure of the complex formed by the membrane type 1-matrix metalloproteinase with the tissue inhibitor of metalloproteinases-2, the soluble progelatinase A receptor. EMBO J. 17, 5238–5248

43. Nivón, L. G., Moretti, R., and Baker, D. (2013) A Pareto-optimal refinement method for protein design scaffolds. PLoS One. 10.1371/JOURNAL.PONE.0059004

44. Mandell, D. J., Coutsias, E. A., and Kortemme, T. (2009) Sub-angstrom accuracy in protein loop reconstruction by robotics-inspired conformational sampling. Nat. Methods. 6, 551–552

45. Krivacic, C., Kundert, K., Pan, X., Pache, R. A., Liu, L., Conchúir, S. O., Jeliazkov, J. R., Gray, J. J., Thompson, M. C., Fraser, J. S., and Kortemme, T. (2022) Accurate positioning of functional residues with robotics-inspired computational protein design. Proc. Natl. Acad. Sci. U. S. A. 119, e2115480119

46. Chevalier, A., Silva, D.-A. A., Rocklin, G. J., Hicks, D. R., Vergara, R., Murapa, P., Bernard, S. M., Zhang, L., Lam, K.-H. H., Yao, G., Bahl, C. D., Miyashita, S.-I. I., Goreshnik, I., Fuller, J. T., Koday, M. T., Jenkins, C. M., Colvin, T., Carter, L., Bohn, A., Bryan, C. M., Fernández-Velasco, D. A., Stewart, L., Dong, M., Huang, X., Jin, R., Wilson, I. A., Fuller, D. H., and Baker, D. (2017) Massively parallel de novo protein design for targeted therapeutics. Nature. 550, 74–79

47. Cao, L., Coventry, B., Goreshnik, I., Huang, B., Sheffler, W., Park, J. S., Jude, K. M., Marković, I., Kadam, R. U., Verschueren, K. H. G., Verstraete, K., Walsh, S. T. R., Bennett, N., Phal, A., Yang, A., Kozodoy, L., DeWitt, M., Picton, L., Miller, L., Strauch, E. M., DeBouver, N. D., Pires, A., Bera, A. K., Halabiya, S., Hammerson, B., Yang, W., Bernard, S., Stewart, L., Wilson, I. A., Ruohola-Baker, H., Schlessinger, J., Lee, S., Savvides, S. N., Garcia, K. C., and Baker, D. (2022) Design of protein-binding proteins from the target structure alone. Nat. 2022 6057910. 605, 551–560

48. Crooks, G. E., Hon, G., Chandonia, J. M., and Brenner, S. E. (2004) WebLogo: A sequence logo generator. Genome Res. 14, 1188–1190

49. Morrison, J. F. (1969) Kinetics of the reversible inhibition of enzyme-catalysed reactions by tight-binding inhibitors. BBA - Enzymol. 185, 269–286

50. Evans, R., O’Neill, M., Pritzel, A., Antropova, N., Senior, A., Green, T., Žídek, A., Bates, R., Blackwell, S., Yim, J., Ronneberger, O., Bodenstein, S., Zielinski, M., Bridgland, A., Potapenko, A., Cowie, A., Tunyasuvunakool, K., Jain, R., Clancy, E., Kohli, P., Jumper, J., and Hassabis, D. (2021) Protein complex prediction with AlphaFold-Multimer. bioRxiv. 10.1101/2021.10.04.463034

51. Jumper, J., Evans, R., Pritzel, A., Green, T., Figurnov, M., Ronneberger, O., Tunyasuvunakool, K., Bates, R., Žídek, A., Potapenko, A., Bridgland, A., Meyer, C., Kohl, S. A. A., Ballard, A. J., Cowie, A., Romera-Paredes, B., Nikolov, S., Jain, R., Adler, J., Back, T., Petersen, S., Reiman, D., Clancy, E., Zielinski, M., Steinegger, M., Pacholska, M., Berghammer, T., Bodenstein, S., Silver, D., Vinyals, O., Senior, A. W., Kavukcuoglu, K., Kohli, P., and Hassabis, D. (2021) Highly accurate protein structure prediction with AlphaFold. Nat. 2021 5967873. 596, 583–589

52. Dauparas, J., Anishchenko, I., Bennett, N., Bai, H., Ragotte, R. J., Milles, L. F., Wicky, B. I. M., Courbet, A., de Haas, R. J., Bethel, N., Leung, P. J. Y., Huddy, T. F., Pellock, S., Tischer, D., Chan, F., Koepnick, B., Nguyen, H., Kang, A., Sankaran, B., Bera, A. K., King, N. P., and Baker, D. (2022) Robust deep learning–based protein sequence design using ProteinMPNN. Science (80-.). 378, 49–56

53. Mehner, C., Hockla, A., Miller, E., Ran, S., Radisky, D. C., and Radisky, E. S. (2014) Tumor cell-produced matrix metalloproteinase 9 (MMP-9) drives malignant progression and metastasis of basal-like triple negative breast cancer. Oncotarget. 5, 2736–2749

54. Akiva, E., Itzhaki, Z., and Margalit, H. (2008) Built-in loops allow versatility in domain-domain interactions: Lessons from self-interacting domains. Proc. Natl. Acad. Sci. U. S. A. 105, 13292–13297

55. Radisky, E. S., Raeeszadeh-Sarmazdeh, M., and Radisky, D. C. (2017) Therapeutic Potential of Matrix Metalloproteinase Inhibition in Breast Cancer. J. Cell. Biochem. 118, 3531–3548

56. Raeeszadeh-Sarmazdeh, M., Coban, M., Mahajan, S., Hockla, A., Sankaran, B., Downey, G. P., Radisky, D. C., and Radisky, E. S. (2022) Engineering of tissue inhibitor of metalloproteinases TIMP-1 for fine discrimination between closely related stromelysins MMP-3 and MMP-10. J. Biol. Chem. 298, 101654

57. Stevens, A. O., and He, Y. (2022) Benchmarking the Accuracy of AlphaFold 2 in Loop Structure Prediction. Biomolecules. 12, 985

58. Erijman, A., Rosenthal, E., and Shifman, J. M. (2014) How structure defines affinity in protein-protein interactions. PLoS One. 9, e110085

59. Kortemme, T., Joachimiak, L. A., Bullock, A. N., Schuler, A. D., Stoddard, B. L., and Baker, D. (2004) Computational redesign of protein-protein interaction specificity. Nat. Struct. Mol. Biol. 11, 371–379

60. Shifman, J. M., and Mayo, S. L. (2003) Exploring the origins of binding specificity through the computational redesign of calmodulin. Proc. Natl. Acad. Sci. U. S. A. 100, 13274–13279

61. Raeeszadeh-Sarmazdeh, M., Greene, K. A., Sankaran, B., Downey, G. P., Radisky, D. C., and Radisky, E. S. (2019) Directed evolution of the metalloproteinase inhibitor TIMP-1 reveals that its N- and C-terminal domains cooperate in matrix metalloproteinase recognition. J. Biol. Chem. 294, 9476–9488

62. Batra, J., Soares, A. S., Mehner, C., and Radisky, E. S. (2013) Matrix Metalloproteinase-10/TIMP-2 Structure and Analyses Define Conserved Core Interactions and Diverse Exosite Interactions in MMP/TIMP Complexes. PLoS One. 8, e75836

63. Mirdita, M., Schütze, K., Moriwaki, Y., Heo, L., Ovchinnikov, S., and Steinegger, M. (2022) ColabFold: making protein folding accessible to all. Nat. Methods 2022 196. 19, 679–682

64. Chao, G., Lau, W. L., Hackel, B. J., Sazinsky, S. L., Lippow, S. M., and Wittrup, K. D. (2006) Isolating and engineering human antibodies using yeast surface display. Nat. Protoc. 1, 755–768

65. Erijman, A., Dantes, A., Bernheim, R., Shifman, J. M., and Peleg, Y. (2011) Transfer-PCR (TPCR): A highway for DNA cloning and protein engineering. J. Struct. Biol. 175, 171–177

66. Benatuil, L., Perez, J. M., Belk, J., and Hsieh, C. M. (2010) An improved yeast transformation method for the generation of very large human antibody libraries. Protein Eng. Des. Sel. 23, 155–159

67. Colby, D. W., Kellogg, B. A., Graff, C. P., Yeung, Y. A., Swers, J. S., and Wittrup, K. D. (2004) Engineering antibody affinity by yeast surface display. Methods Enzymol. 388, 348–358

68. Whitehead, T. A., Chevalier, A., Song, Y., Dreyfus, C., Fleishman, S. J., De Mattos, C., Myers, C. A., Kamisetty, H., Blair, P., Wilson, I. A., and Baker, D. (2012) Optimization of affinity, specificity and function of designed influenza inhibitors using deep sequencing. Nat. Biotechnol. 30, 543–548

69. Wrenbeck, E. E., Faber, M. S., and Whitehead, T. A. (2017) Deep sequencing methods for protein engineering and design. Curr. Opin. Struct. Biol. 45, 36–44

70. Gaspar, J. M. (2018) NGmerge: Merging paired-end reads via novel empirically-derived models of sequencing errors. BMC Bioinformatics. 19, 1–9

71. Rubin, A. F., Gelman, H., Lucas, N., Bajjalieh, S. M., Papenfuss, A. T., Speed, T. P., and Fowler, D. M. (2017) A statistical framework for analyzing deep mutational scanning data. Genome Biol. 18, 1–15

72. Love, M. I., Huber, W., and Anders, S. (2014) Moderated estimation of fold change and dispersion for RNA-seq data with DESeq2. Genome Biol. 15, 1–21

73. Batra, J., Robinson, J., Soares, A. S., Fields, A. P., Radisky, D. C., and Radisky, E. S. (2012) Matrix metalloproteinase-10 (MMP-10) interaction with tissue inhibitors of metalloproteinases TIMP-1 and TIMP-2: Binding studies and crystal structure. J. Biol. Chem. 287, 15935–15946

74. Suzuki, K., Kan, C. C., Hung, W., Gehring, M. R., Brew, K., and Nagase, H. (1998) Expression of human pro-matrix metalloproteinase 3 that lacks the N-terminal 34 residues in Escherichia coli autoactivation and interaction with tissue inhibitor of metalloproteinase 1 (TIMP-1). Biol. Chem. 379, 185–191

75. Mehner, C., Hockla, A., Miller, E., Ran, S., Radisky, D. C., Radisky, E. S., Mehner, C., Hockla, A., Miller, E., Ran, S., Radisky, D. C., and Radisky, E. S. (2014) Tumor cell-produced matrix metalloproteinase 9 (MMP-9) drives malignant progression and metastasis of basal-like triple negative breast cancer. Oncotarget. 5, 2736–2749

