## Supplementary Information for "A broad matrix metalloproteinase inhibitor with designed loop extension exhibits ultrahigh specificity for MMP-14"

**Supplementary information of the manuscript by Bonadio et al. “****A broad matrix metalloproteinase inhibitor with designed loop extension exhibits ultrahigh specificity for MMP-14”**

**Table S1.** Sequences of the seven designs.

|  | **67** | **67A** | **67B** | **67C** | **67D** | **67E** | **68** |
| --- | --- | --- | --- | --- | --- | --- | --- |
| **Des4** | **A** | **M** | **E** | **D** | **V** | **W** | **G** |
| **Des1** | **D** | **D** | **T** | **T** | **I** | **I** | **S** |
| **Des2** | **D** | **T** | **D** | **G** | **I** | **W** | **G** |
| **Des22** | **D** | **T** | **D** | **G** | **V** | **F** | **G** |
| **Des14** | **A** | **T** | **N** | **D** | **I** | **T** | **D** |
| **Des15** | **A** | **N** | **S** | **G** | **I** | **I** | **D** |
| **Des10** | **M** | **V** | **S** | **T** | **M** | **M** | **G** |

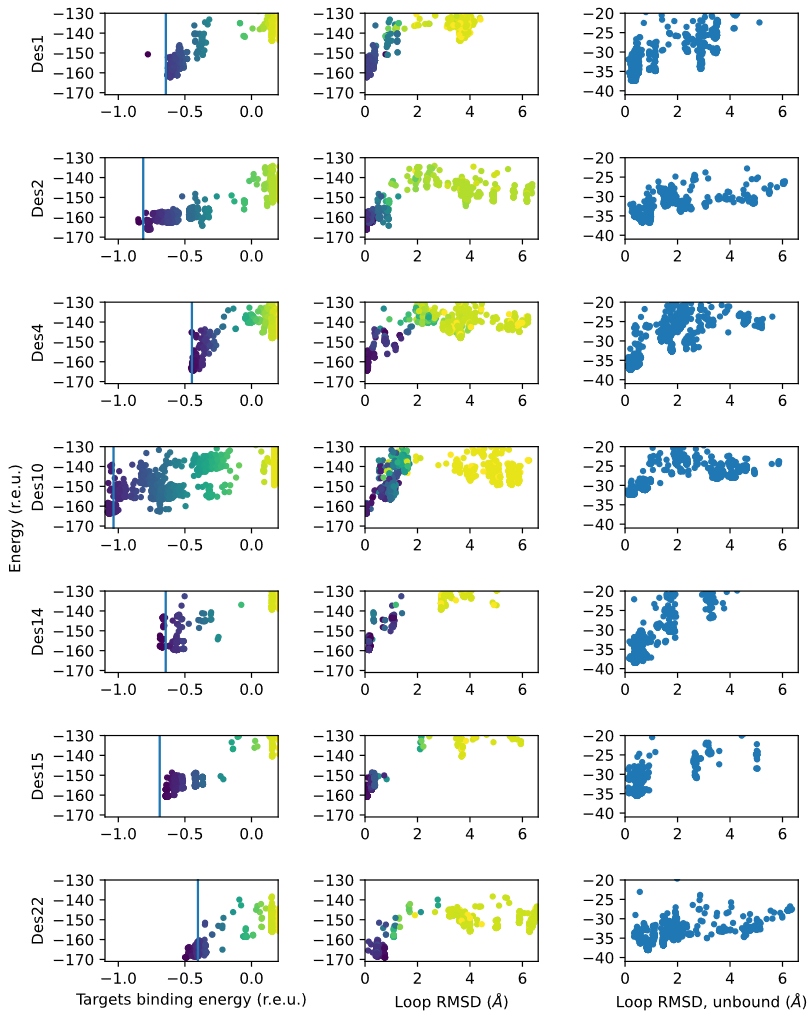
**Figure S1. Selection of designs by loop modeling, using the sequences of the designed loops**. Left column, Energy vs target-residues binding energy. The vertical line marks the energy in the designed original loop model. The target-residues binding energy is also color-coded from purple (low) to yellow (high). Central column, Energy vs loop RMSD plot, in the bound setup (MMP-14/N-TIMP2). The target-residues binding energy is color-coded from purple to yellow. All seven designs showed one low energy conformation having 0-1 Å RMSD from the intended design and with the indented target binding energy (dark purple). Right column, Energy vs loop RMDS plot in the unbound state, showing that the intended conformation is preorganized in most designs.

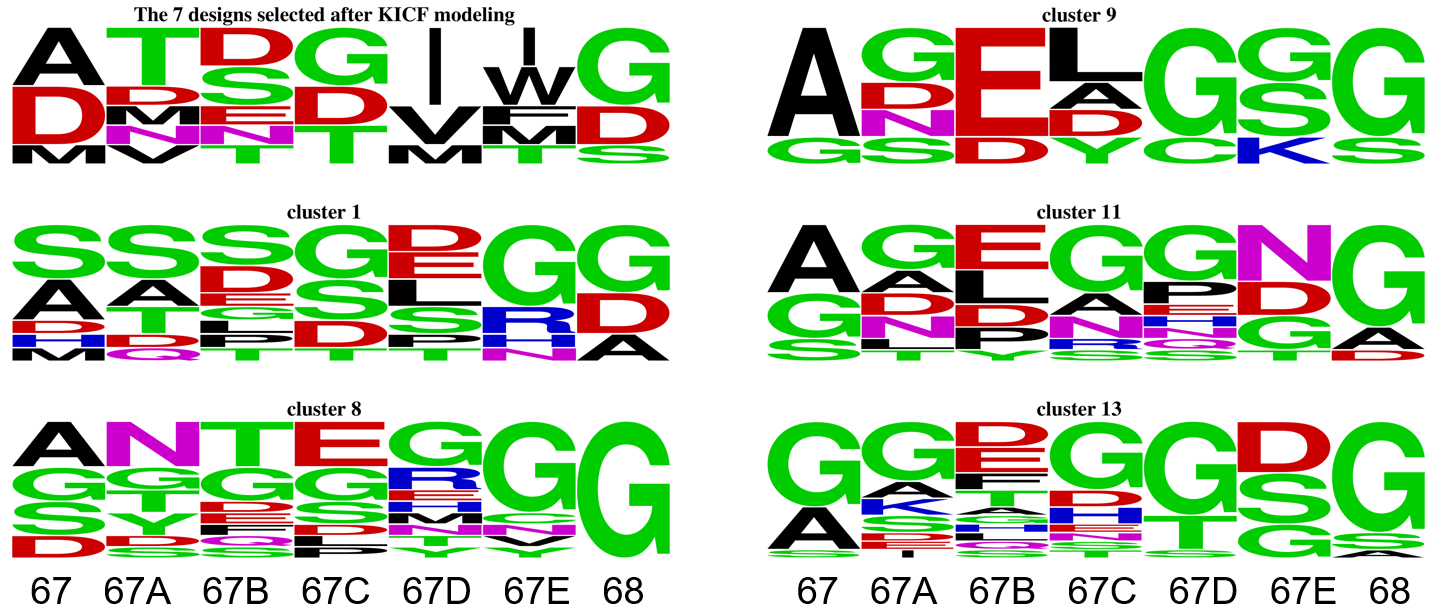

**Figure S2. Sequence logos of loop clusters with conformations reaching the non-conserved regions**. Designs from a non-constrained loop design run with Rosetta Remodel were clustered and 26 clusters were visually inspected selecting the clusters reaching the non-conserved region on MMPs. Cluster 1 reaches E248, while the other clusters reach G210 and P207. The sequence logo of the 7 designs selected after FKIC is also shown. Position 67 and 68 are at the ends of the designed loop and therefore more spatially restrained in possible loop conformations. The logos show low variability among clusters, particularly in the 7 designs selected after FKIC (contacting G210 and P207) and cluster 1 (contacting E248). Sequence logos were done using WebLogo (1).

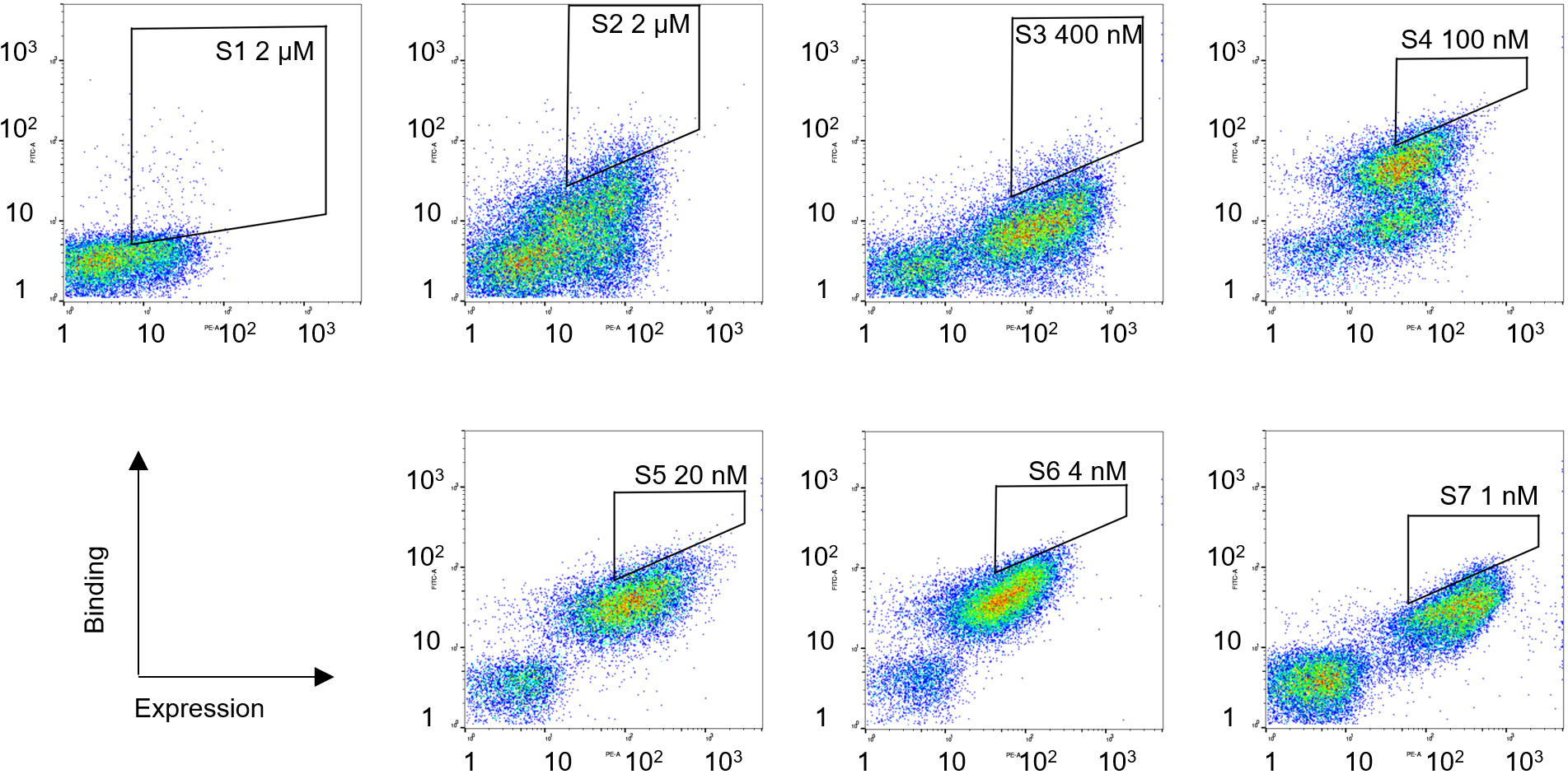

**Figure S3. The affinity maturation FACS sorts.** Gates are labeled with the name of the output sorted population indicating the sorting cycle, and the MMP concentration used.

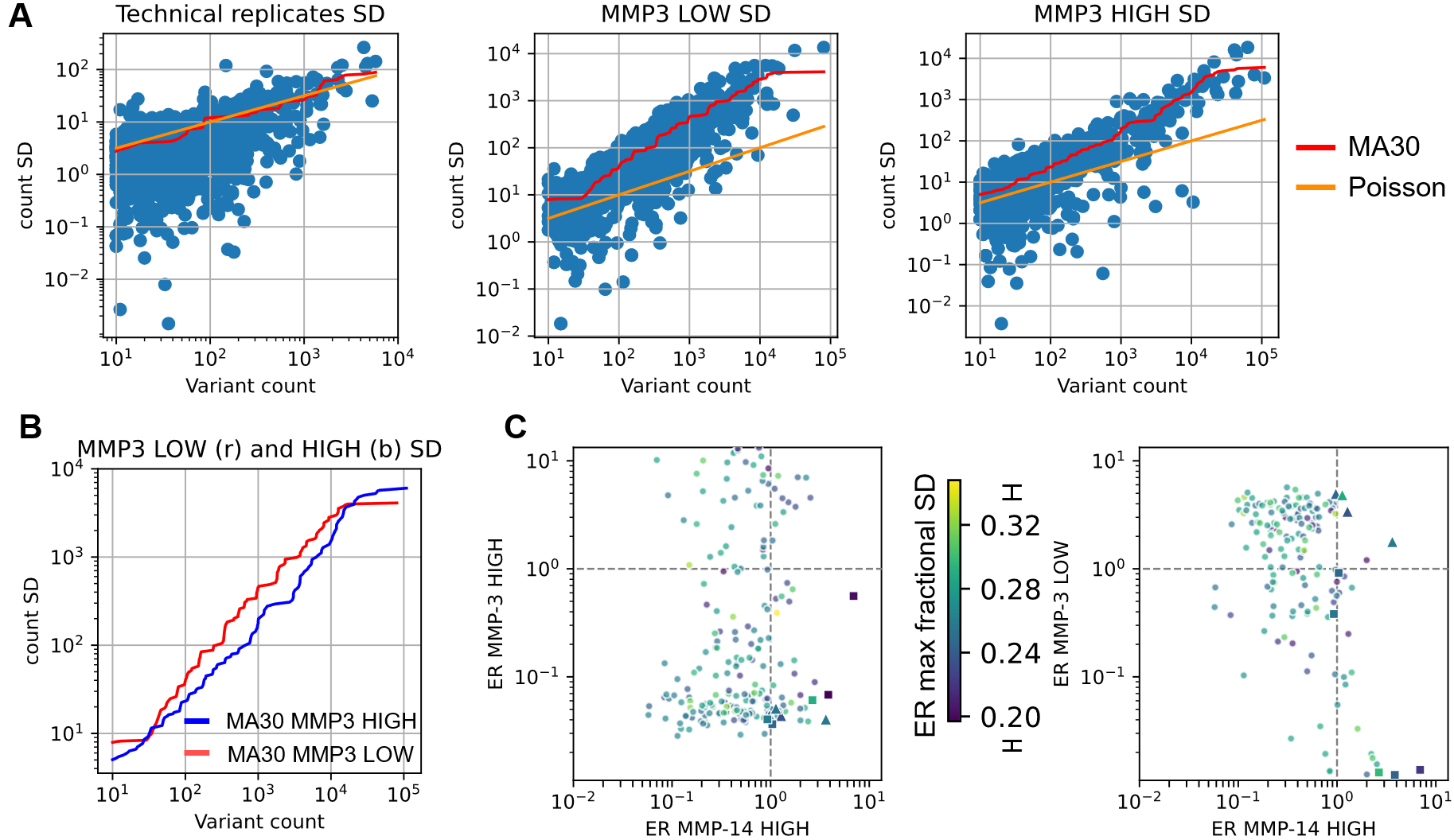

**Figure S4. Calculation of variant count standard deviations.** A, the moving average method (MA) of the duplicate count standard deviations (SDs) along the average variant counts (red), or simple Poisson model SD = sqrt(n), where n is the count (orange). 30-window centered MA of the frequency standard deviations along the variant frequency was calculated, and then transformed for the variant count. Left, technical replicates in which the NGS runs were repeated with identical samples. Center and Right, independent biological duplicates (repeated FACS and NSG) in the two different setups S6_MMP-3,LOW_ and S6_MMP-3,HIGH_ showed in Fig. 2B. The Poisson model and MA are in good agreement for the technical replicates but show big deviations for the independent duplicates as previously reported with more complex statistical models (2, 3). B, calculated SD using MA from the two different setups showed in A. center and right, are compared showing overall robustness. C, ER for non-target MMP-3 vs ER for the target MMP-14 is shown analogous to Fig 2C. Color represents the maximum fractional SD of the two ERs for MMP-14 and MMP-3, propagated from the MA30 S6_MMP-3,HIGH_. At the color bar 0.2 and 0.32 ticks, corresponding error bars are indicated, in the figures scale.

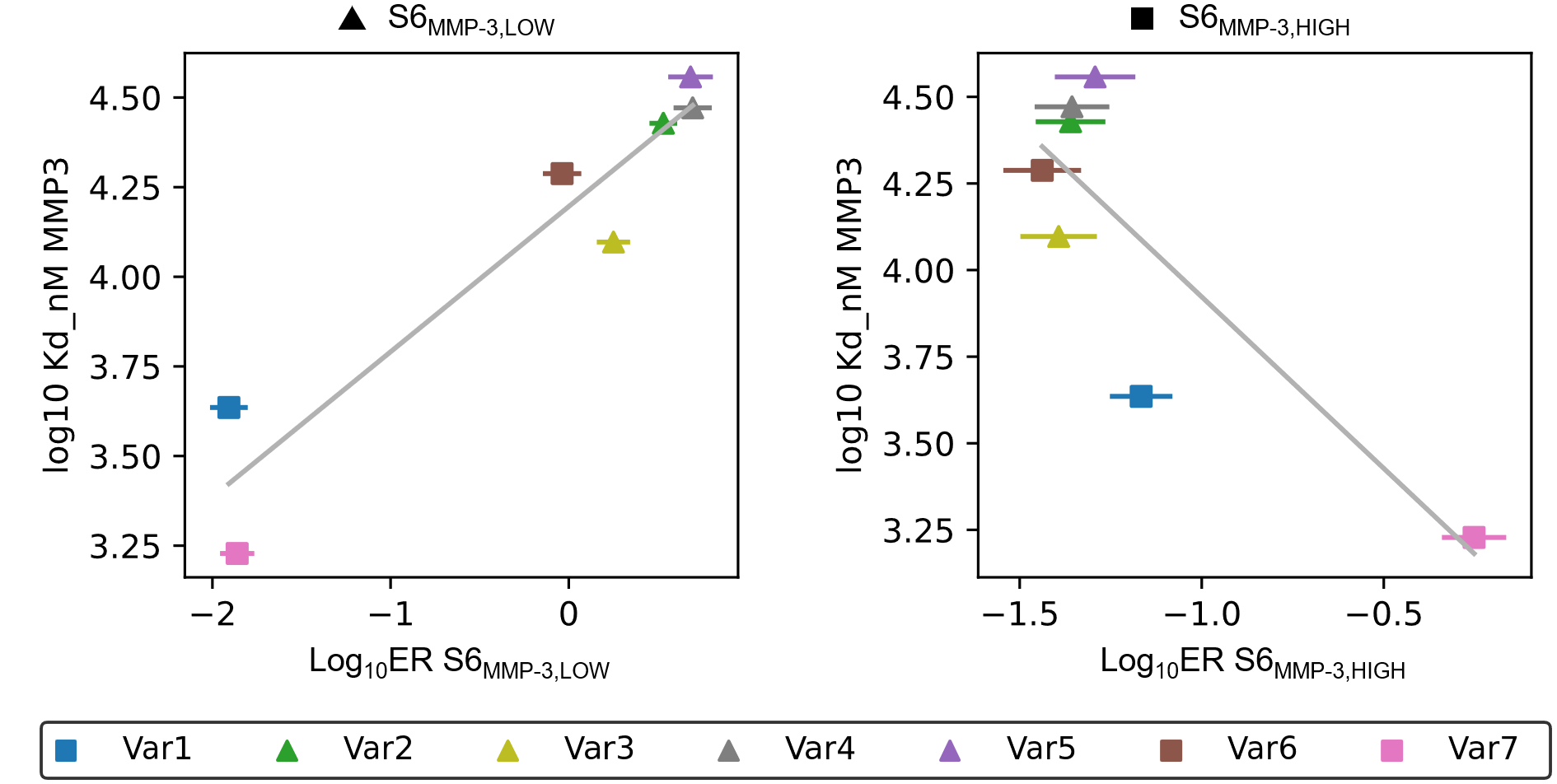

**Figure** **S5. Correlation between ER and Kds measured by YSD titration.** The type of MMP-3 sort shown in figure 2C from which the variant was selected is indicated for each of the 7 variants tested by yeast titration, S6_MMP-3,LOW_ with triangles and S6_MMP-3,HIGH_ with squares. The ERs correlate with K_D_s measured by YSD, especially for the S6_MMP3,LOW_ sort (Spearman’s correlation r=0.89, p=0.01).

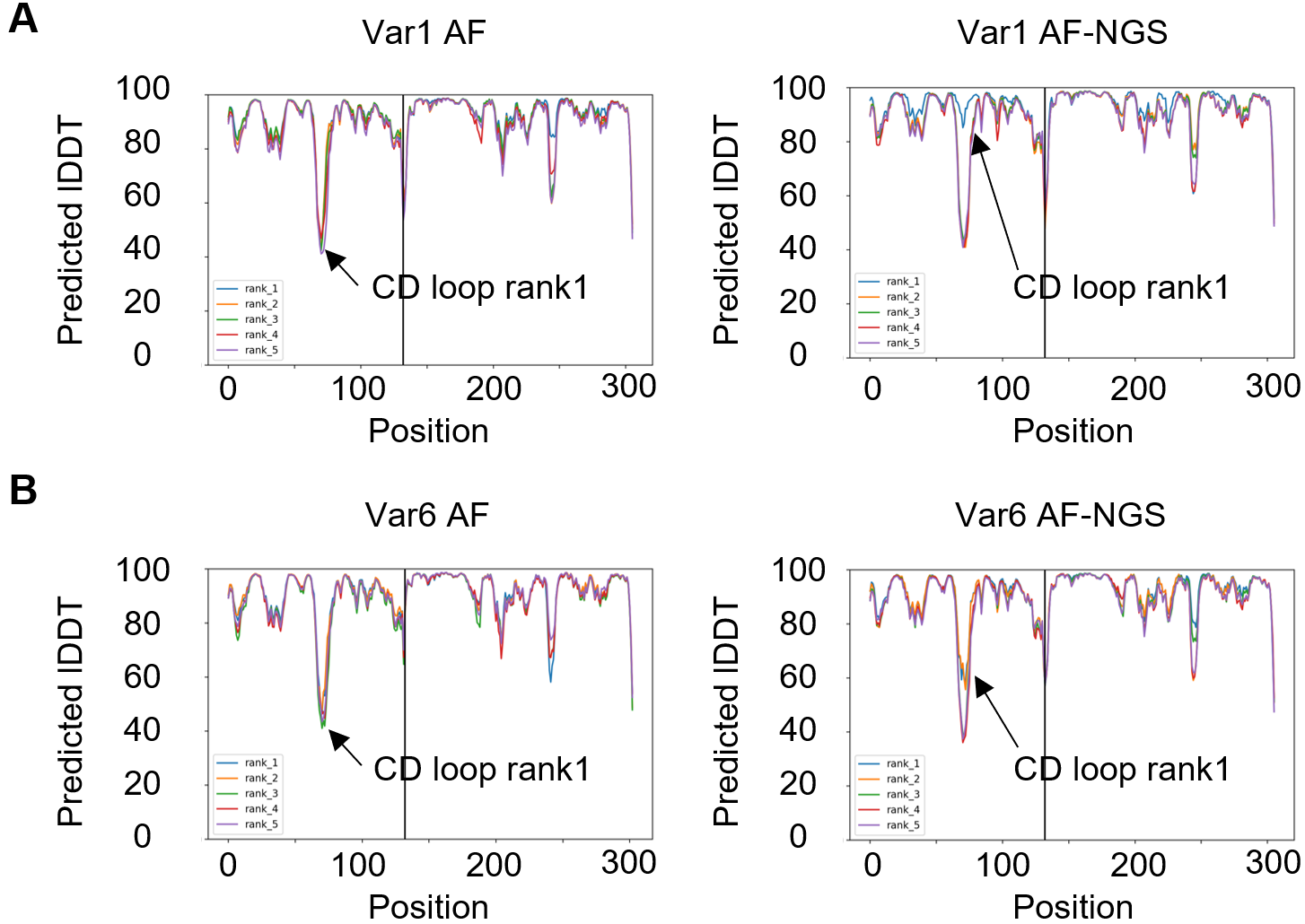

**Figure S6.** **AlphaFold CD loop accuracy is greatly increased with the addition of NGS sequences in the MSA.** A, variant Var1 model accuracy in the engineered loop region is increased by adding the sequences found in cluster 2, to which Var1 belongs to the AlphaFold input MSA. B, an increase in accuracy is also observed for variant Var6.

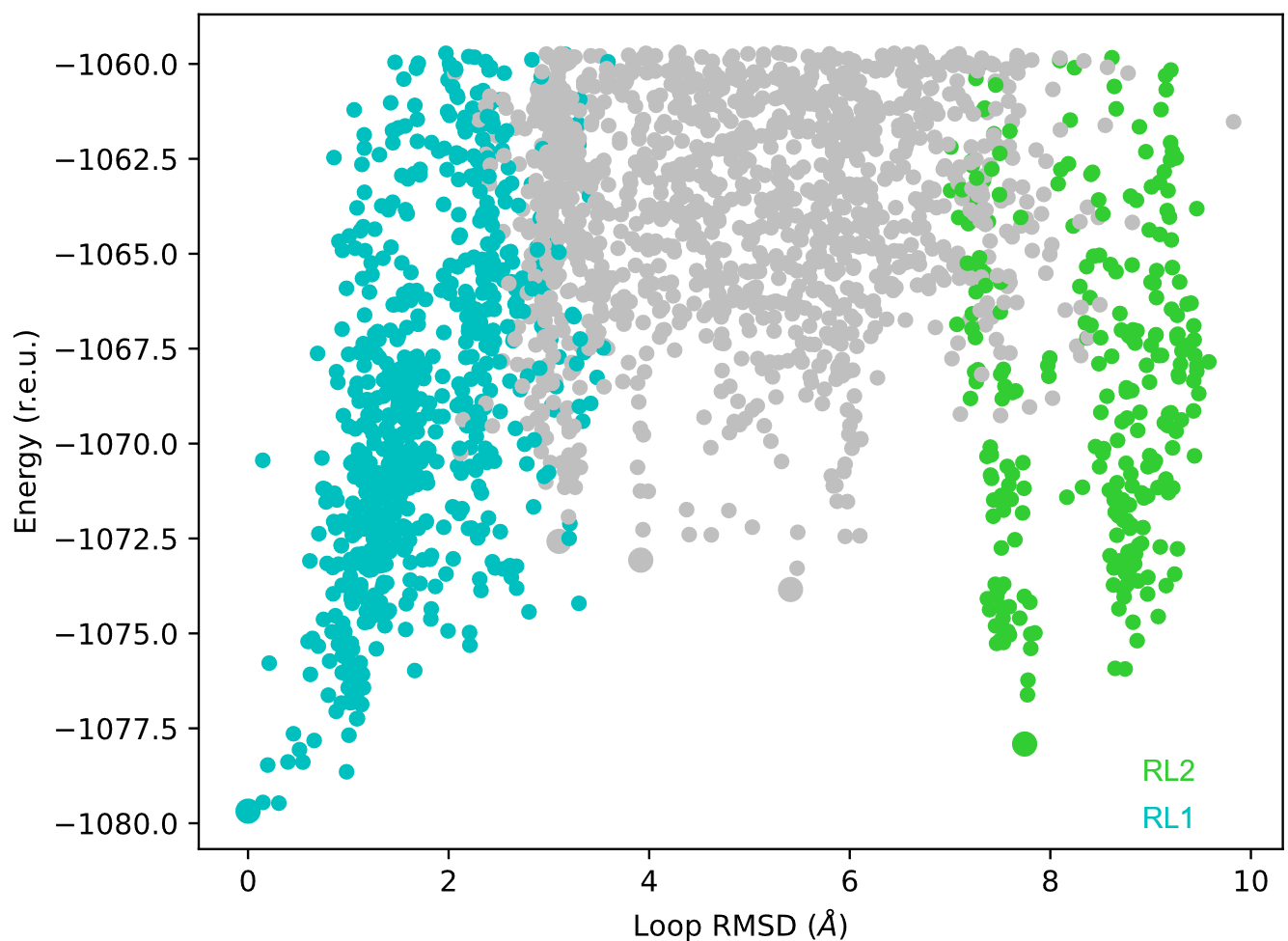
**Figure S7. Conformational sampling of the N-TIMP2 variant Var1 engineered loop.** Energy vs loop RMSD from the lowest-energy model found in the modeling procedure. Only models with energy within the lowest 20 r.e.u. are shown. The two lowest energy clusters (4 Å cutoff) are colored, showing the RL1 model (cyan) and the RL2 model (green). The lowest energy models from the lowest 5 clusters are shown in bigger circles.

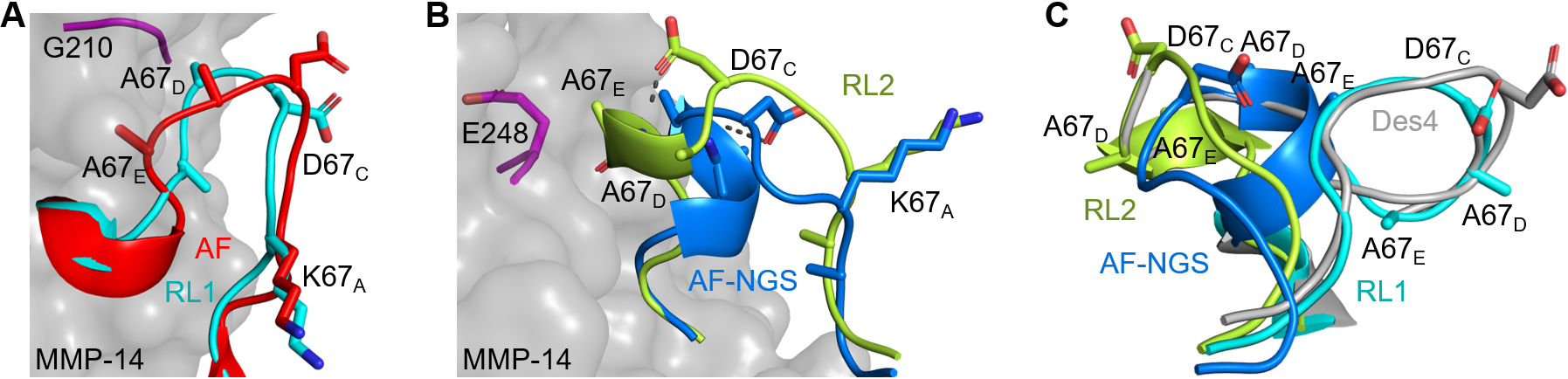

**Figure S8. Structural models of variant Var1 in complex with MMP-14.** All models share a topologically similar conformation, with K67_A_ anchoring the loop to the rest of the N-TIMP2, D67_C_ at the tip of the loop, and A67_D_ and A67_E_ at the distal binding interface. A, The AF (red) and the RL1 (cyan) models. The models are similar in overall conformation, especially at the N- and C-termini and in the positioning of K67_A_, D67_C_, and A67_D_. B, The AF-NGS (blue) and the RL2 (light-green) models. The models are similar in overall conformation, especially at the N- and C-termini, and in the positioning of K67_A_. C, Superposition of AF-NGS (blue), RL1 (cyan), and RL2(green) and Des4 (gray) loop models. All models exhibit local similarities. The fragment composed of D67_C_, A67_D_, A67_E_ was found in the 7 designs shown in figure 1D and superposes well with RL1, AF-NGS and RL2 (RMSD of 0.08 Å, 0.04 Å, 0.09 Å respectively). The fragment backbone from Des4 superposed to each model is shown.

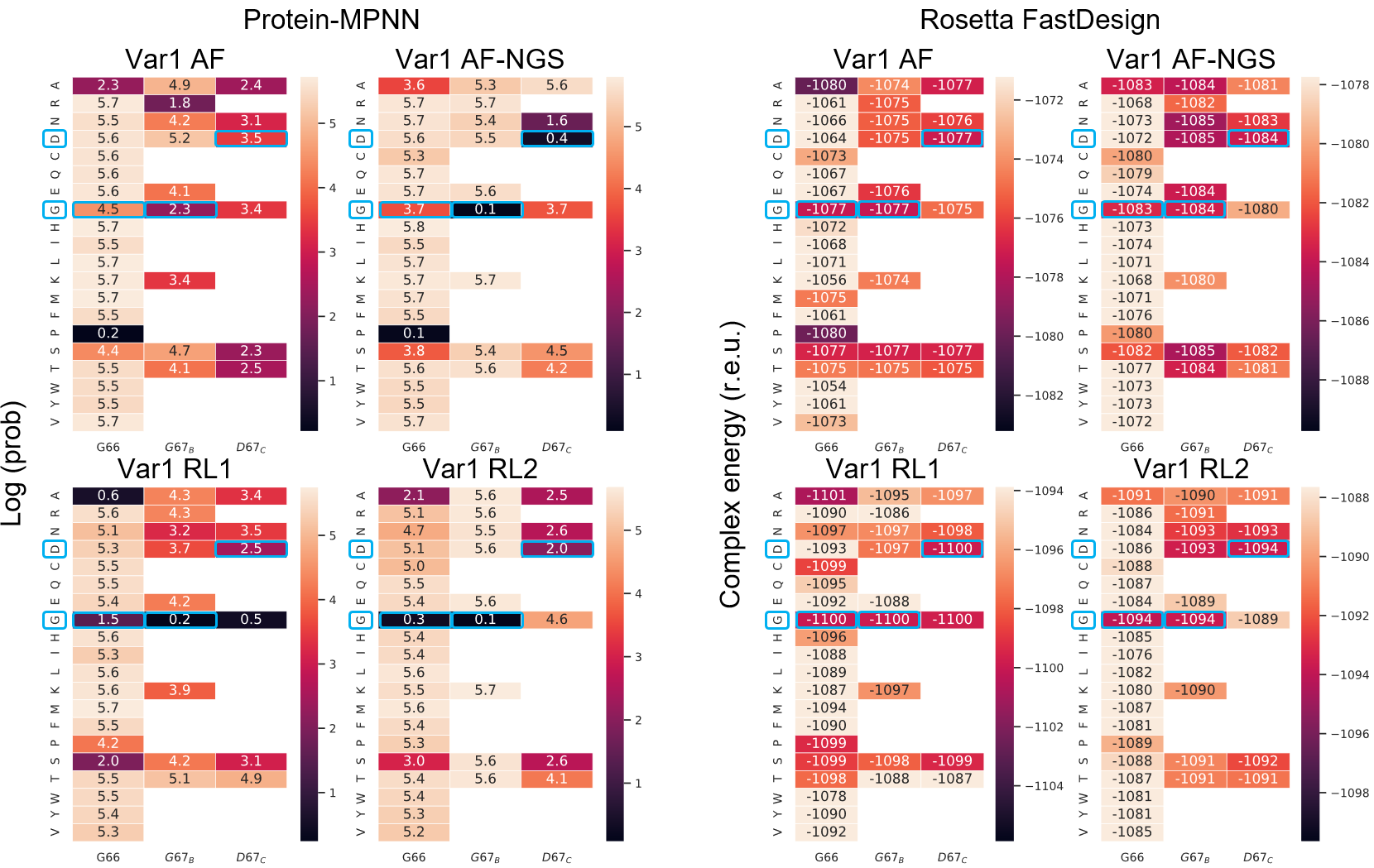

**Figure S9. Sequence design recapitulates the amino acid conservation for the structures of the AF-NGS and RL2 loop models for variant Var1.** The AF model, the AF-NGS model, the RL1, and the RL2 of variant Var1 were sequence designed by predicting the positional probability of the full loop using the protein-MPNN approach (left) and the computational single position mutational scanning in Rosetta FastDesign (right). Amino acids not present in the library are not shown. Cluster 2 has a highly conserved G66/ G67_B_/D67_C_ motif, and Var1 AF-NGS shows the highest conservation at G67_B_/D67_C_ in protein-MPNN, with the same position predicted as favorable in Rosetta fast design. We noted that at position 66 an unexpected proline is predicted by protein-MPNN but not by Rosetta fast design. RL2 model and to a lesser extent RL1 show high conservation at G66 ad G67_B_, while D67_C_ is predicted to be less conserved. In contrast, AF shows the lowest conservation of these three residues.

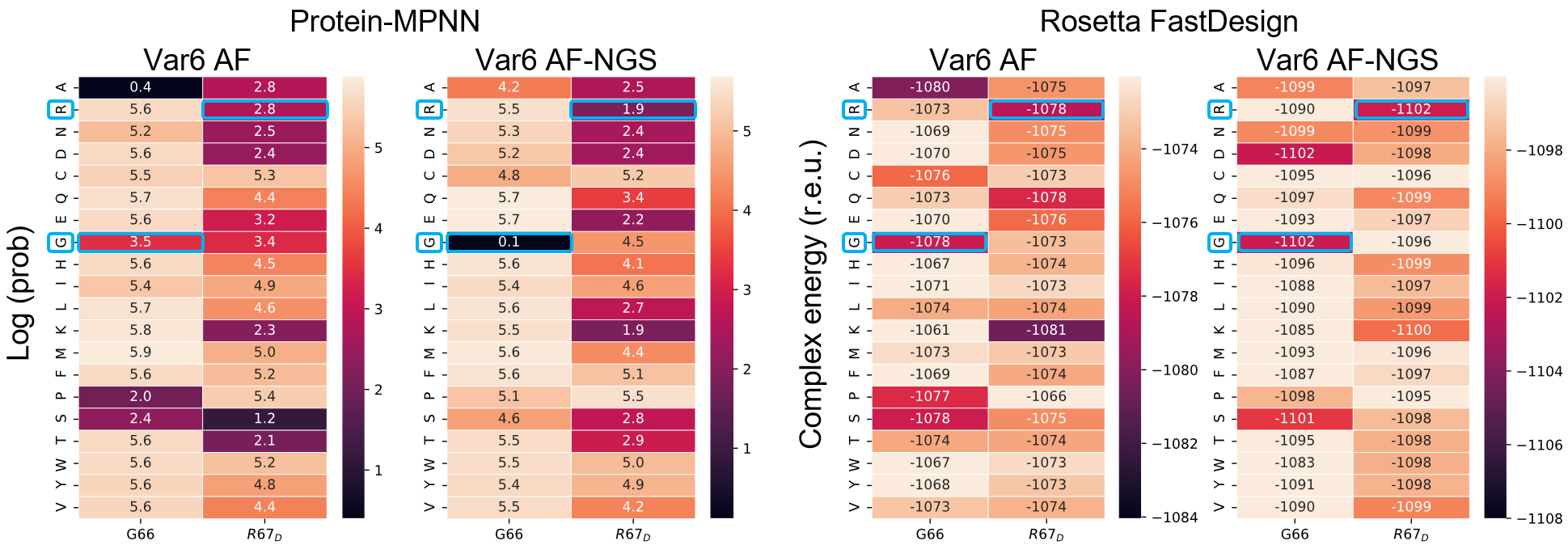

**Figure S10. Sequence design recapitulates the amino acid conservation for the structure of the AF-NGS loop model for variant Var6**. Analogous to Supplementary Fig. S9 but for Var6. The AF and the AF-NGS model of variant Var6 were redesigned by predicting the position probability of the full loop with protein-MPNN (left) and by computational mutational scanning in Rosetta fast design (right). Amino acids not belonging to the library are shown white. Var6 belongs to Cluster 1, which has the conserved motif G66/R67_D._ AF-NGS shows conservation at G66/R67_D_, while the AF model allows amino acids not found in the cluster profile.

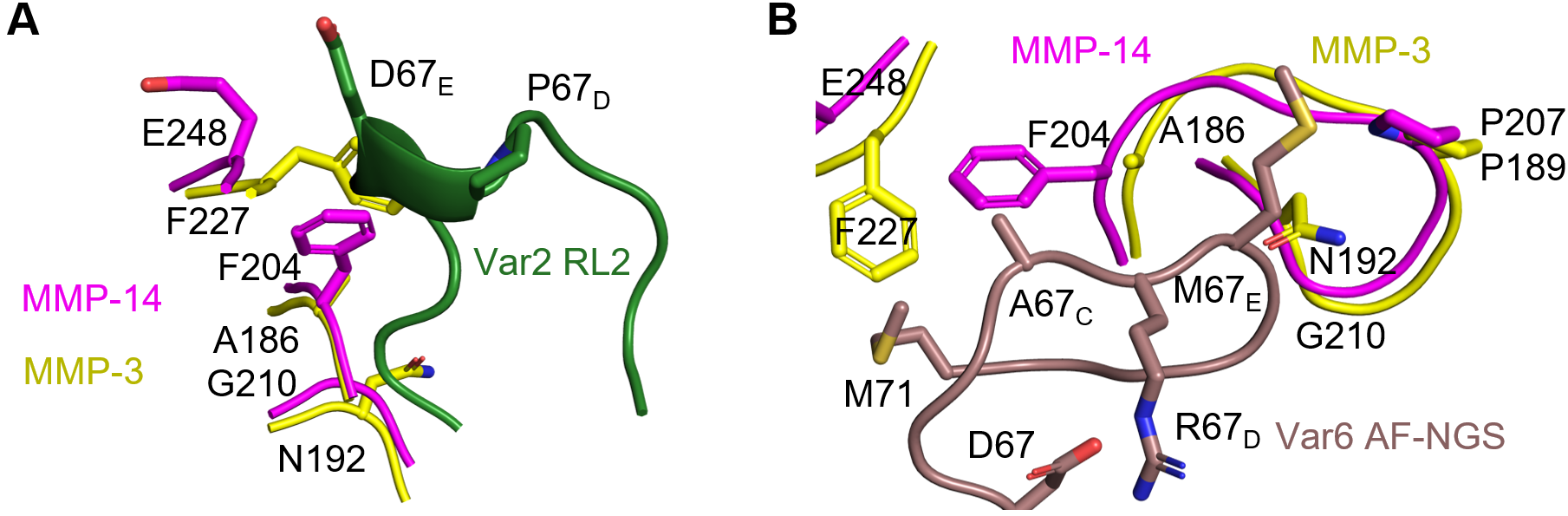

**Figure S11.** **Structural models of variant Var2 and Var6 in complex with MMP-14 and MMP-3.** A, the Rosetta top-ranking model for Var2, very similar to the Var1 RL2 model. An apparent clash with F227 on MMP-3 is analogous to the one found for Var1 in Fig. 4C. B, the AF-NGS top ranking model for Var6. The conserved R67_D_ stabilizes the loop with a salt bridge to D67. The loop is further stabilized with 9 intramolecular H-bonds and 1 intramolecular H-bond. A67_C_, M67_E_ and M71 form hydrophobic interactions. The differences between MMP-14 and MMP-3 environment, especially at E248/F227, G210/N192, F204/A186, might weaken those hydrophobic interactions and be responsible for observed Var6 specificity. N192 on MMP-3 is also creating a small apparent clash with the M67_E_ backbone, which might further contribute to specificity.

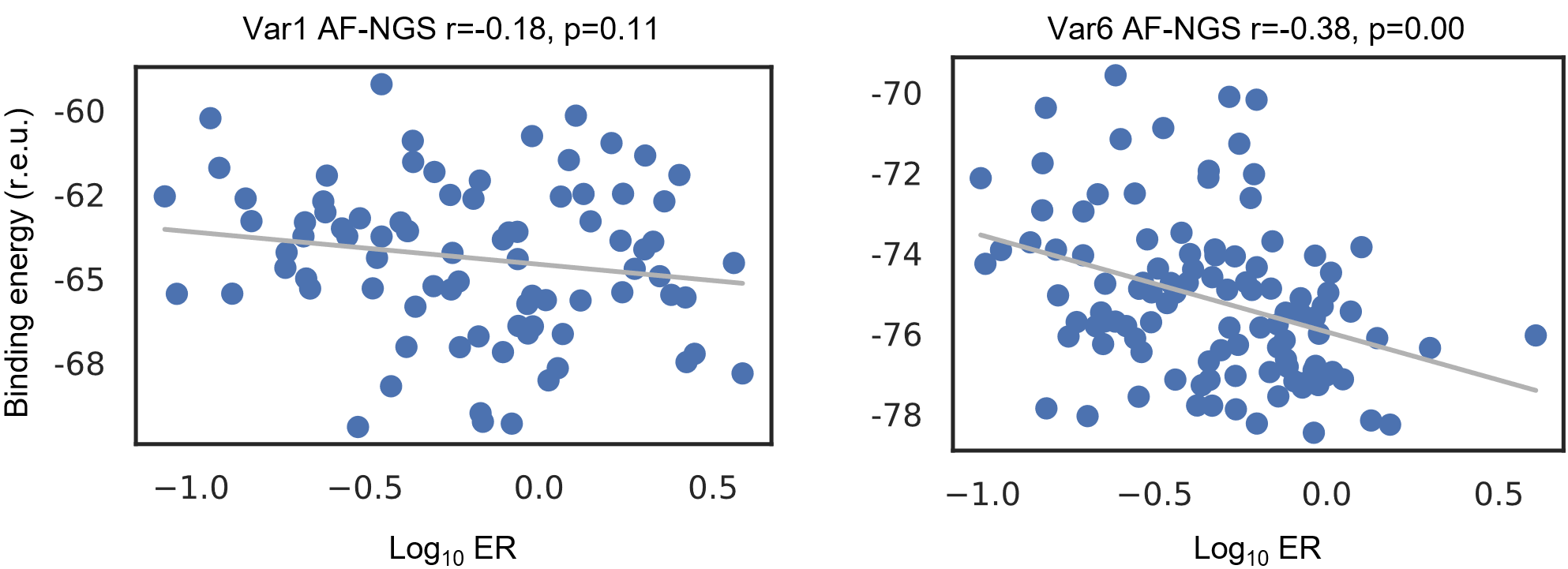

**Figure S12. Correlation between Rosetta binding energy and MMP-14 ERs for AF-NGS models**. The AF-NGS model for Var1/MMP-14 (left) or Var6/MMP14 (right) were used to thread the sequences of their respective clusters, cluster 2 for Var1 and cluster 1 for Var6 (Fig. 2D). The binding energy calculated with Rosetta Interface Analyzer(4) shows negative correlation with ER of the variant in the sort. A weak correlation could be expected since variants typically differ by 3-5 mutations located in the loop, and mutations stabilizing the loop in the binding conformation entropically contribute to ΔG_bind_ since the loop is likely to have some flexibility, but this contribution is not accounted for in the simple method used here, which only calculated the interaction energy by separating the complex. Spearman’s correlation is shown in plot titles.

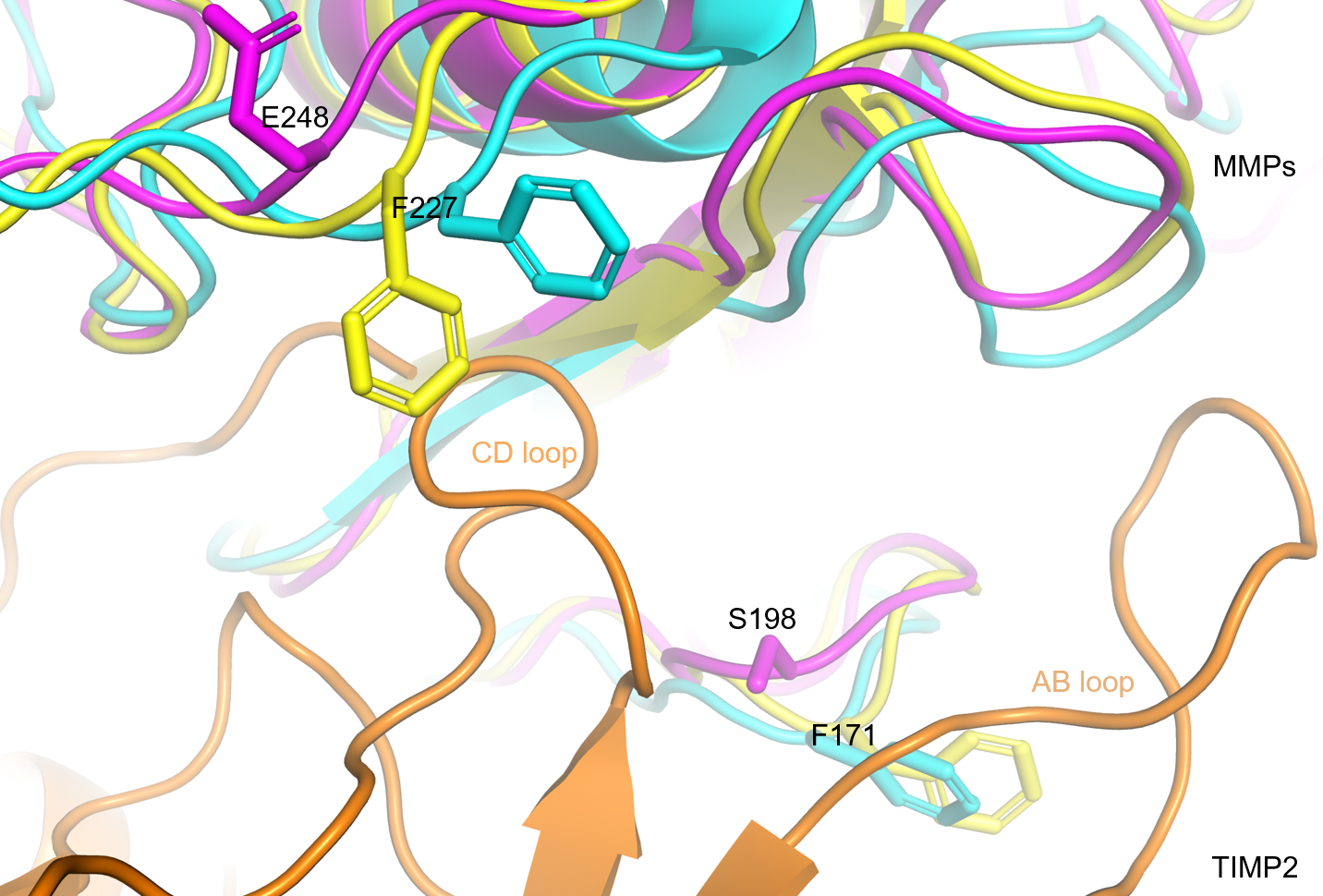

**Figure S13. The two structural models for the N-TIMP2/MMP-3 complex**. Models are viewed superposed on N-TIMP2. Cyan: MMP-3 from the N-TIMP2/MMP-3 model based on the alignment of MMP-3 (6mav) to the TIMP2/MMP-10 structure (4ilw), yellow: MMP-3 from the AlphaFold N-TIMP2/MMP-3 model, magenta and orange: MMP-14 and N-TIMP2 from the TIMP2/MMP-14 structure (1buv). F171 in MMP-3 protrudes towards the N-TIMP2 AB loop. This phenylalanine is not present in MMP-14, which has S198 instead. In the structural alignment model of N-TIMP2/MMP-3, F171 “pushes” the N-TIMP2 AB loop, tilting N-TIMP2, similar to the TIMP2/MMP10 structure in 4ilw. This accentuates the difference in the location of another phenylalanine F227 in MMP-3, compared to E248 in MMP-14. This difference is less prominent in the AlphaFold model of the N-TIMP2/MMP-3 complex because here the MMP-3 backbone at F171 rearranges to accommodate the N-TIMP2 AB loop, while the rest of the backbone maintains a mode of binding more similar to TIMP2/MMP-14 in 1buv, and therefore there is less tilting of TIMP2.

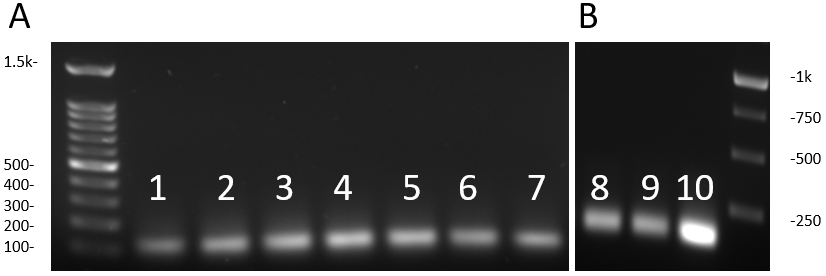

**Figure S14.** A, The mixed-bases ultramer was annealed and extended to a partially complementary primer. The reaction was carried on at 72 degrees for different time intervals: 1: 1’, 2: 2’, 3: 10’, 4: 30’, 5: 60’, 6: 90’ 7: 90’. B, Amplification of the library at various primer concentrations: 8: 0.5 µM, 9: 1 µM, 10: 2 µM.

References

1.	Crooks, G. E., Hon, G., Chandonia, J. M., and Brenner, S. E. (2004) WebLogo: A sequence logo generator. *Genome Res.* **14**, 1188–1190

2.	Love, M. I., Huber, W., and Anders, S. (2014) Moderated estimation of fold change and dispersion for RNA-seq data with DESeq2. *Genome Biol.* **15**, 1–21

3.	Rubin, A. F., Gelman, H., Lucas, N., Bajjalieh, S. M., Papenfuss, A. T., Speed, T. P., and Fowler, D. M. (2017) A statistical framework for analyzing deep mutational scanning data. *Genome Biol.* **18**, 1–15

4.	Benjamin Stranges, P., and Kuhlman, B. (2013) A comparison of successful and failed protein interface designs highlights the challenges of designing buried hydrogen bonds. *Protein Sci.* **22**, 74–82
